# Trial-by-trial modulation of express visuomotor responses induced by symbolic or barely detectable cues

**DOI:** 10.1101/2021.01.29.428908

**Authors:** Samuele Contemori, Gerald E. Loeb, Brian D. Corneil, Guy Wallis, Timothy J. Carroll

## Abstract

Human cerebral cortex can produce visuomotor responses that are modulated by contextual and task-specific constraints. However, the distributed cortical network for visuomotor transformations limits the minimal response time of that pathway. Notably, humans can generate express visuomotor responses that are inflexibly tuned to the target location and occur 80-120ms from stimulus presentation (stimulus-locked responses, SLRs). This suggests a subcortical pathway for visuomotor transformations involving the superior colliculus and its downstream reticulo-spinal projections. Here we investigated whether cognitive expectations can modulate the SLR. In one experiment, we recorded surface EMG from shoulder muscles as participants reached toward a visual target whose location was unpredictable in control conditions, and partially predictable in cue conditions by extrapolating a symbolic cue (75% validity). Valid symbolic cues led to faster and larger SLRs than control conditions; invalid symbolic cues produced slower and smaller SLRs than control conditions. This is consistent with a cortical top-down modulation of the putative subcortical SLR-network. In a second experiment, we presented high-contrast targets in isolation (control) or ~24ms after low-contrast stimuli, which could appear at the same (valid cue) or opposite (invalid cue) location as the target, and with equal probability (50% cue validity). We observed faster SLRs than control with the valid low-contrast cues, whereas the invalid cues led to the opposite results. These findings may reflect exogenous priming mechanisms of the SLR network, potentially evolving subcortically via the superior colliculus. Overall, our results support both top-down and bottom-up modulations of the putative subcortical SLR network in humans.

**NEW & NOTEWORTHY:** Express visuomotor responses in humans appear to reflect subcortical sensorimotor transformation of visual inputs, potentially conveyed via the tecto-reticulo-spinal pathway. Here we show that the express responses are influenced both by symbolic and barely detectable spatial cues about stimulus location. The symbolic cue-induced effects suggest cortical top-down modulation of the putative subcortical visuomotor network. The effects of barely detectable cues may reflect exogenous priming mechanisms of the tecto-reticulo-spinal pathway.

## INTRODUCTION

Extraction of information about the surrounding environment is crucial to guide motor behaviour in everyday life and sport contexts, but also to react to threatening events for survival. In higher vertebrates, the availability of a cerebral cortex enables extrapolation of surrounding sensory cues and generation of expectations about probable future events. These expectations can facilitate the transformation of expected sensory information into motor responses, thus reducing the reaction time (RT; see for review Posner 2016; van Ede et al. 2012).

Humans are capable of generating extremely rapid (*express*) responses to visual stimuli (Pruszynski et al. 2010). As opposed to the so-called volitional muscle response, the initiation time of these early EMG responses does not co-vary with the movement onset time and is consistently within 80-120ms after stimulus presentation (Pruszynski et al. 2010; Wood et al. 2015). Therefore, these express visuomotor responses have been called stimulus-locked responses (SLRs; see Contemori et al. 2020 for discussion of appropriate nomenclature). Furthermore, the SLR is always directed toward the stimulus location irrespective of whether the task requires to move toward (pro-reach) or against (anti-reach) the stimulus (Gu et al. 2016), or to withhold the movement (Atsma et al. 2018). It is worth noting that the short-latency and inflexible characteristics of SLRs are also properties of express saccades, which are generated subcortically via the superior colliculus and its downstream projections to the reticular formation (Dorris et al. 1997; Pare and Munoz 1996; Fischer and Boch 1993). Therefore, the SLR may also result from subcortical sensorimotor transformation of visual inputs through the tecto-reticulo pathway and its downstream projections to the spinal motoneurons and interneurons (see for review Corneil and Munoz 2014).

The occurrence of express saccades increases as a function of collicular *pre-target* activity level (Dorris et al. 1997; Dorris et al. 2002), probably via a direct influence on collicular *target-related* response amplitude. For instance, cueing the target with a prior (~50ms) stimulus at the same location (i.e. valid cue) has been shown to prime the pre-target activity of superior colliculus neurons and amplify the ensuing target-related response (Fecteau et al. 2004). This facilitates both rapid initiation of saccades (Fecteau et al. 2004) and neck muscle SLRs (Corneil et al. 2008) as compared with no-cued and invalidly cued targets, a phenomenon known as *attention capture* (for review see Klein 2000; Corneil and Munoz 2014). These observations suggest that target-directed visuomotor behaviours are modulated as a function of pre-target sensory events and their influence on visuomotor networks, including the superior colliculus and its downstream reticulo-spinal circuits.

In the first experiment, we tested the hypothesis that pre-target signals affording cognitive expectations about the location of approaching targets can modify the SLR expression. Therefore, we employed a pre-target cue whose information depended on its perceived orientation rather than its location, thus requiring cognitive extrapolation. In the second experiment, we used a different target-cueing paradigm to study the influence of barely detectable visual events on visuomotor behaviour, and tested the hypothesis that SLRs are participant to bottom-up priming effects. The purpose of this paper was to delineate the influence of symbolic and barely detectable visual cues on express visuomotor behaviour. This would provide evidence about the influence of both top-down and bottom-up neural modulation mechanisms of the SLR and its putative underlying subcortical network, including the superior colliculus. The findings may contribute to our understanding of the neural mechanisms underlying express visuomotor behaviour in humans.

## MATERIALS AND METHODS

### Participants

Sixteen adults participated in the first experiment (14 males, 2 females; mean age: 31.6 years, SD: 6.9), and twelve of them also completed the second experiment (11 males, 1 female; mean age: 31.3 years, SD: 6.0). All participants were right-handed, had normal or corrected-to-normal vision, and reported no current neurological, or musculoskeletal disorders. They provided informed consent and were free to withdraw from the experiment at any time. All procedures were approved by the University of Queensland Medical Research Ethics Committee (Brisbane, Australia) and conformed to the Declaration of Helsinki.

### Apparatus

The apparatus used for this study has been previously described by Contemori et al. (2020). Briefly, the participants performed target-directed reaching movements with their dominant hand via shoulder extension (right ward), or flexion (left ward), movements in the transverse plane. Because muscle pre-activation has proven effective to facilitate SLR expression (Gu et al. 2016; Contemori et al. 2020), a constant lateral load of ~5N was applied in the direction of transverse shoulder extension via a weight and pulley system. This increased the baseline activity of shoulder transverse flexor muscles, including the clavicular head of pectoralis major muscle.

All stimuli were created in Matlab using the Psychophysics toolbox (Brainard 1997; Pelli 1997), and were displayed on a LCD monitor with a 120Hz refresh rate (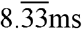/refresh cycle) positioned ~57cm in front of the participants. For the first experiment, the target was a full and filled black circle of ~2dva in diameter presented against a light grey background. This created a high target-to-background contrast (luminance: black target, ~0.3 cd/m^2^; grey background, ~137 cd/m^2^) which has been shown to enhance SLR expression (Wood et al. 2015). Conversely, in the second experiment we used high-contrast (~0.3 cd/m^2^) and low-contrast targets, which were both full filled circles of ~2dva in diameter. For each participant, the low-contrast target luminance was customized to visual acuity (see below for details). On average, the low-contrast stimulus luminance was ~119.7cd/m^2^. The luminance was measured with a colorimeter (Cambridge Research System ColorCAL MKII). A photodiode was attached to the left bottom corner of the monitor to detect a secondary light that was presented coincidentally with the time of appearance of the real target. This allowed us to index the time point at which the stimulus was physically detectable, thus avoiding uncertainties in software execution and raster scanning of the monitor.

### Experimental design

#### Experiment 1: symbolic cue

This experiment was designed to investigate the influence of cognitive expectations on express visuomotor responses. The participants were instructed to reach as fast as possible toward a visual target that appeared as a brief flash of a complete circle, features that facilitate SLRs (Contemori et al. 2020; Kozak et al. 2019). The target location was unpredictable or partially predictable from the orientation of a symbolic arrow-shaped cue (Figure 1). The stimuli were presented via an *emerging target* paradigm (Figure 1) that has proven effective for facilitating the SLR expression in more than 80% of paricipants tested with surface EMG electrodes (Contemori et al. 2020), and that was motivated by preceding SLR (Kozak et al. 2020) and oculomotor studies (for review see Fiehler et al. 2019). To start the trial, the participants aligned their right hand and gaze for one second on a fixation spot (“+” sign) located in the centre of the screen and below the visual barrier (~9dva of fixation-target eccentricity). After the fixation period, the central fixation spot could remain unchanged (neutral cue, control condition) or change to an arrow pointing to the future location of the target (valid cue, 75% of cue trials) or in the wrong direction (invalid cue, 25% of cue trials). Note that the physical position of the cue was irrelevant with respect to the future target locations. At ~700ms after the cue presentation, the target dropped at constant velocity (~35dva/s) toward the visual barrier for ~160ms, and always re-emerged (‘go’ signal) below it after ~640ms from the onset of its movement (i.e. predictably timed stimulus). Therefore, the target was occluded by the barrier for ~480ms and re-emerged after ~1.34s from the cue presentation (Figure 1). We decided to use a cue-target onset asynchrony (CTOA) of more than 1 second in order to ensure unambiguous cognitive extrapolation of the arrow orientation. Note that the temporal events timings have been adjusted by rounding the values to the nearest ten milliseconds (full monitor scanning occurred every ^—^ms, see previous section).

**Figure 1:**
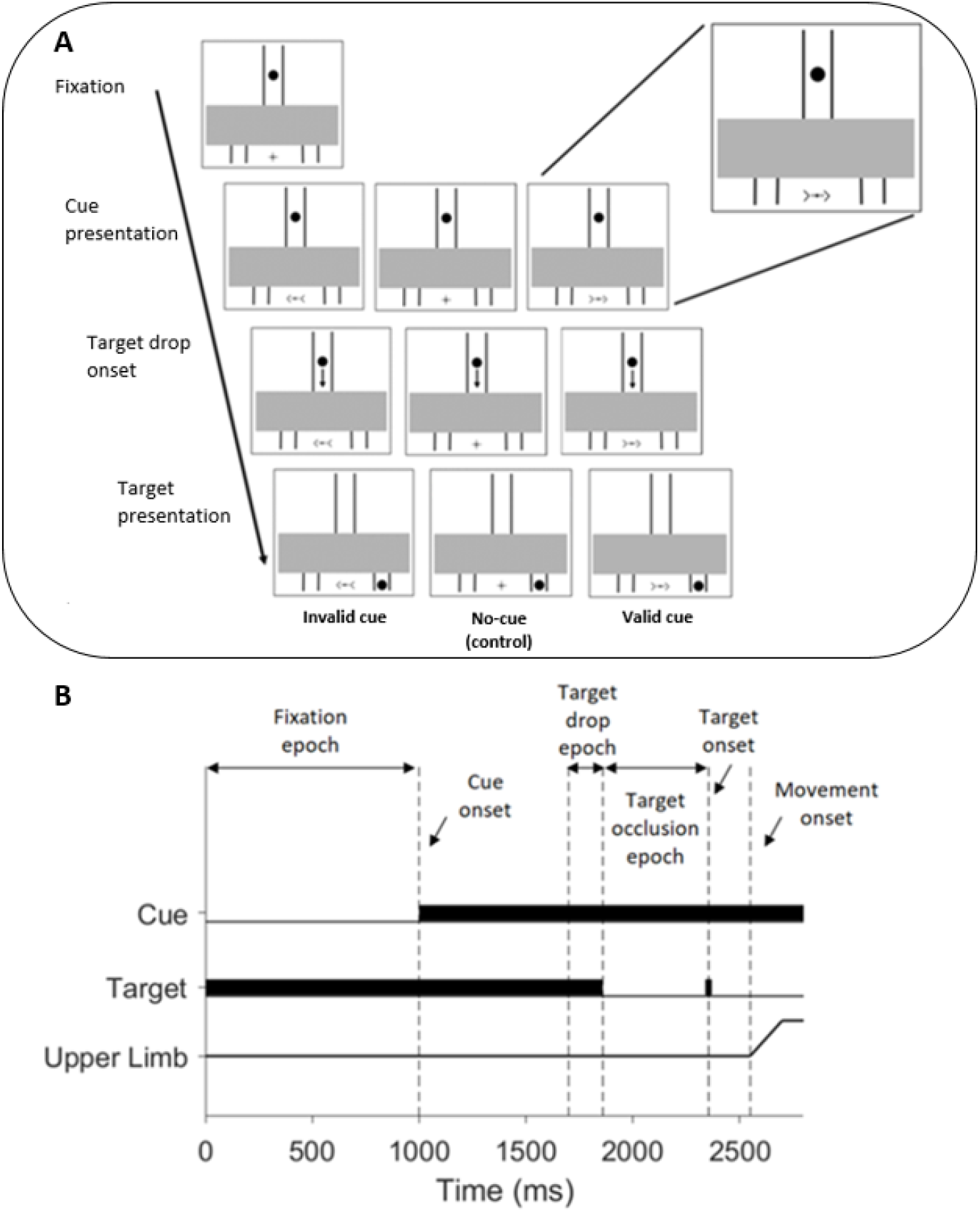
(A) Timeline of no-cue (control), valid and invalid cue conditions of the first experiment. A zoomed view of the symbolic arrow-shaped cue is shown in the top right corner. In these examples, the target appears to the right so the right inset panels show a valid cue trial, whereas the left inset panels show an invalid cue trial. (B) Schematic diagram of temporal events in the cue conditions. After one second of fixation, the central cross bar for fixation remained unchanged in the control condition whereas it was substituted by an arrow cue pointing toward the exact future location of the target (valid cue, 75% of cue trials) or in the wrong direction (invalid cue, 25% of cue trials). After ~700ms from cue presentation, the target started dropping from the stem of the track at constant velocity of ~35dva/s until it passed behind the barrier (occlusion epoch) for ~480ms, and re-appeared underneath it at ~640ms from the onset of its movement. The target appeared transiently by making one single flash of ~8ms of duration.

On each trial, gaze-on-fixation was checked on-line with an EyeLink 1000 plus tower-mounted eye tracker device (SR Research Ltd., Ontario, Canada), at a sampling rate of 1000 Hz. If the fixation requirement was not met, participants received an error message and the trial was repeated. Each participant completed 10 blocks of 72 reaches/block (36 for each direction), with each block consisting of 46 valid, 16 invalid and 10 neutral cues, randomly intermingled.

#### Experiment 2: low-contrast cue

In this experiment, we aimed to investigate whether the SLR is modified by spatially cueing the target location with barely detectable cues. For each participant, we initially set the target-luminance threshold for stimulus detection as a function of visual acuity via an adaptive (staircase) procedure (Kindom and Prins 2016). The task was the same as the control conditions in the first experiment, but the circle started dropping immediately after 1 second of fixation (Figure 2) and the luminance of the target flashing underneath the barrier was changed trial-by-trial depending on preceding response. Specifically, we generated an array of twenty-two logarithmic scaled steps of luminance ranging from high-contrast target luminance (~0.3 cd/m^2^) to background luminance (~137 cd/m^2^). The participants were required to reach toward the first target flash they perceived below the barrier as soon as possible, and to guess the target location by moving arbitrarily right or left if nothing was perceived. If the movement direction was correct (see below), then the target luminance was made dimmer (i.e. closer to background colour) by selecting the next luminance level in the array (i.e. one step up). By contrast, if the movement was incorrect the target luminance was made four times darker than the last flashed target (i.e. four steps down in the array - this only happened when the target was at least five steps dimmer than the high-contrast target). No-movement trials were also classified as incorrect movements. Further, random jumps of target luminance were used in order to avoid trial-by-trial dependencies (Kindom and Prins 2016). The staircase procedure was terminated after ten reversals (i.e. wrong reach made after a correct response) of the target luminance, which occurred on average after ~65 trials. The final low-contrast stimulus used in the second experiment (Figure 2) had the average luminance used in the 10 trials before the last reversal, corresponding to correct stimulus detectability on ~80% of presentation as per the “1up/4down” staircase approach (Kindom and Prins 2016).

**Figure 2:**
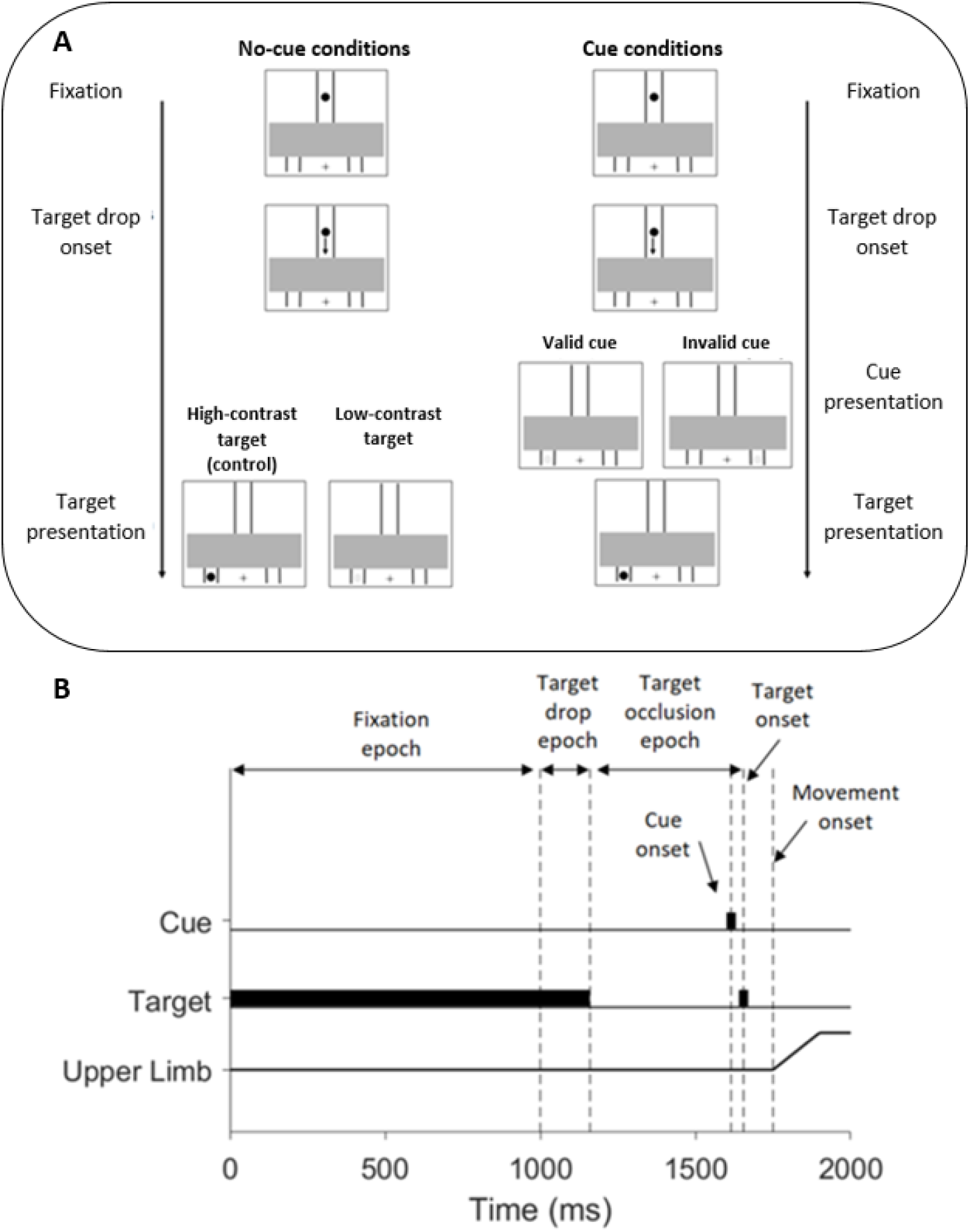
(A) Timeline of high-contrast (control condition) target, low-contrast (dim grey dot) target, valid, and invalid cue conditions of the second experiment. In these examples, the high-contrast target appears to the left, so the valid cue condition is satisfied when the low-contrast stimulus (dim grey dot) appears to the left, whereas it appears to the right in the invalid cue condition. The low-contrast cue appeared with equal probability at the same (valid cue) or opposite (invalid cue) location of the ensuing high-contrast target (i.e. 50% cue validity). (B) Schematic diagram of temporal events in the cue conditions. After one second of fixation at the central cross bar, the target started dropping from the stem of the track at constant velocity of ~35dva/s until it passed behind the barrier (occlusion epoch) for ~480ms. The low-contrast cue appeared after ~616ms from the trial start and stayed on for ~8ms. The high-contrast target re-emerged transiently (one single flash of ~8ms of duration) underneath the barrier after ~640ms form the trial start. Therefore, the temporal gap between the low-contrast cue and the high-contrast target was ~24ms.

For the main experiment, we used four unique target conditions: (I) high-contrast (control) target appearing alone underneath the barrier; (II) low-contrast targets appearing alone underneath the barrier; (III) low-contrast cue appearing at the same location of the high-contrast target (valid cue); (IV) low-contrast cue appearing at the opposite location of the high-contrast target (invalid cue). In the cue conditions, the high-contrast target was validly or invalidly cued with equal probability (i.e. 50% cue validity). The low-contrast cue appeared three frames (~24ms) before the high-contrast target, by making a single flash of ~8ms of duration (Figure 2). Importantly, the dim luminance, short CTOA and irrelevant validity (50%) of the low-contrast cues were designed to minimize the involvement of cortical networks in cue processing. Moreover, the brief ~24ms CTOA was chosen in order to avoid *inhibition of return*, a phenomenon known to reverse the advantaging and disadvantaging effects that are otherwise induced by validly and invalidly cueing a target, respectively (for review see Klein 2000). On each trial, the target that dropped toward the barrier was always a full and filled black circle, thus making impossible for the participants to predict the target condition from trial context. The participants were instructed to reach as fast as possible toward the first perceived target flash underneath the barrier, and to guess the target location by reaching arbitrarily right or left if no stimulus was detected. They completed 10 blocks of 64 reaches/block, with each block consisting of 16 trials of each of the 4 different target conditions, randomly intermingled.

### Data recording

Surface EMG (sEMG) activity was recorded from the clavicular head of the right pectoralis muscle (PMch) and the posterior head of the right deltoid muscle (PD), with double-differential surface electrodes (Delsys Inc. Bagnoli-8 system, Boston, MA, USA). The quality of the signal was checked with an oscilloscope before the start of recording. The sEMG signals were amplified by 1000, filtered with a 20-450Hz bandwidth filter by the native ‘Delsys Bagnoli-8 Main Amplifier Unit’, and full-wave rectified after digitization without further filtering. Arm motion was monitored by a three-axis accelerometer (Dytran Instruments, Chatsworth, CA; Contemori et al., 2020). The sEMG and kinematic data were sampled at 2 kHz with a 16-bit analog-digital converter (USB-6343-BNC DAQ device, National Instruments, Austin, TX, USA). Data synchronization was guaranteed by starting the recording of the entire data-set at the frame at which the target started moving toward the barrier.

Reaction time (RT) was monitored by running a cumulative sum analysis (Basseville and Nikiforov 1993) on the acceleration signal, as described in Contemori et al., 2020. In order to minimize the occurrence of anticipatory responses, we monitored the RT online and sent an error message if the participants moved before the target onset time or responded in less than 130ms from target presentation (~3 trials/block). This RT cut-off was adopted because 130ms has been recently shown to be the critical time to prepare a target-directed response (Haith et al. 2016). Furthermore, the initiation of a movement requires agonist muscles activation and antagonist muscles inhibition in order to generate enough net joint torque to overcome limb inertia and produce angular acceleration at the joint. If a target-directed movement occurs faster than 130ms, the potential short-latency sEMG response occurring in the SLR epoch (i.e. 80-120ms from target onset) could be contaminated by an anticipatory voluntary response. This would make impossible to distinguish the SLR from the muscle activity that is time-locked with the voluntary movement initiation. To further reduce this risk, we adopted a more conservative RT cut-off for offline data analysis, by excluding trials with RT<140ms (~7% of the trials).

The accelerometer signal also allowed us to identify correct and wrong responses. Specifically, we searched for the first peak/valley of acceleration subsequent to the RT index in order to define the initial movement direction. We then compared the movement direction with the target location. If the target location did not correspond with the movement direction, the trial was classified as incorrect and discarded (see results). This analysis was run online for the staircase procedure adopted in the second experiment to customize the low-contrast target luminance on each participant visual acuity (see above).

### Data analysis

#### Indexing the presence, timing and magnitude of SLRs

The presence of a candidate SLR was identified with a time-series receiver operator characteristic (ROC) analysis. This analysis allowed us to index the point in time at which the location of the target could be discriminated (discrimination time, DT) from the sEMG trace (Pruszynski et al. 2010). For every muscle sample and tested condition not showing anticipatory activity (for details see Contemori et al. 2020), we sorted the correct trials according to RT and subdivided the sEMG trials into two equally-sized trial sets by doing a median split on the RT data (Figure 3A and D). We then ran separate ROC analyses on the fastest 50% (*fast* trial set) and the slowest 50% (*slow* trial set) of the trials to extrapolate the area under the ROC curve (AUC). The AUC values range from 0 to 1, where a value of 0.5 indicates chance discrimination, whereas a value of 1 or 0 indicates perfectly correct or incorrect discrimination, respectively. We set the thresholds for discrimination at 0.65 (Figure 3B and E); this criterion exceeds the 95% confidence intervals of data randomly shuffled with a bootstrap procedure. The time of earliest discrimination was defined as the time after stimulus onset at which the AUC overcame the defined threshold, and remained above that threshold level for at least 15ms. The candidate SLR was considered only if both fast and slow trial discrimination times were within 80-120ms after target presentation (Gu et al. 2016; Contemori et al. 2020). Further, we associated the fast and slow DTs with the average RT of fast and slow data sets (Wood et al. 2015), and we fitted a line to the data to test if the DT did not co-vary with the RT (i.e. line slope >67.5°, Figure 3C; for further details see Contemori et al., 2020). In this case, we ran the ROC analysis on all trials to extrapolate the *all-trials* set DT (Figure 3E). Finally, we defined the SLR initiation time by running a two-pieces “DogLeg” linear regression analysis (Carroll et al. 2019; Pruszynski et al. 2008) recently adopted by Contemori et al. (2020) to index the point in time at which the time-series ROC curve begins to deviate positively toward the 0.65 discrimination threshold (Figure 3E). Importantly, this analysis allowed us to extrapolate the EMG response initiation time regardless of the slope of the ROC curve as it deviated toward the discrimination threshold (Contemori et al. 2020).

**Figure 3:**
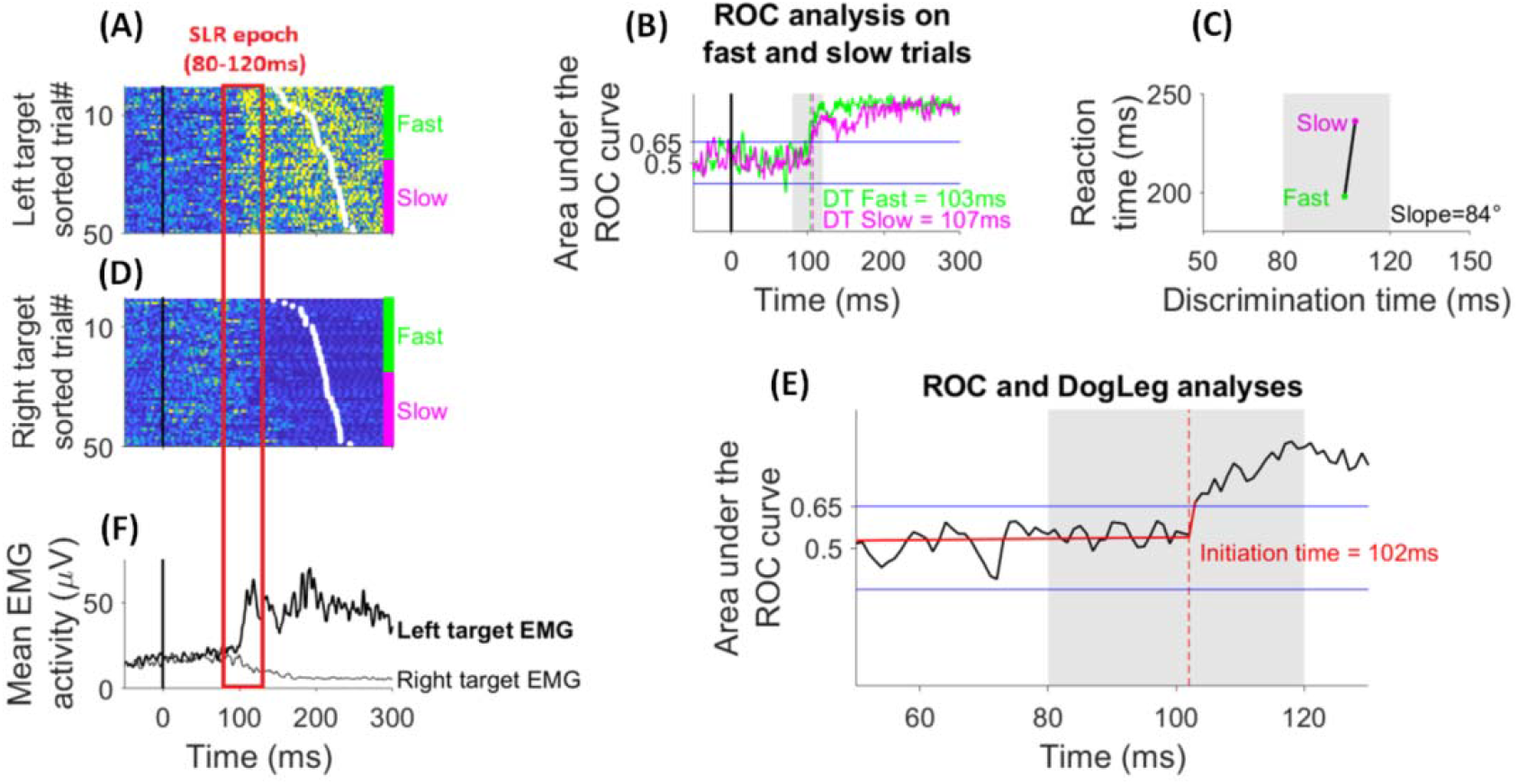
Exemplar sEMG activity from the clavicular head of pectoralis major of a participant who exhibited an SLR in the control condition of the first experiment (participant 8, table 1). The muscle acts as agonist and antagonist for (A) left and (D) right targets, respectively. Rasters of rectified surface sEMG activity from individual trials are shown (darker yellow colours indicate greater sEMG activity; panel A and D) as are the traces of the (F) mean sEMG activity (thick line = left target EMG; thin line = right target EMG). Data are aligned on visual target presentation (solid black vertical line at time 0) and sorted according to reaction time (white dots within the rasters).The unfilled red rectangle indicates the time window in which an SLR is expected (80-120ms from target onset).The SLR appears as a column of either rapid muscle activation (A) or inhibition (D) time-locked to the stimulus onset in both the fastest 50% (green bar) and the slowest 50% (magenta bar) of the trials. (B) ROC analysis panel showing the point in time at which the target location can be discriminated (discrimination time - DT) from muscle activity for the fast (green line) and slow (magenta line) sets of trials. The DT is identified by the first time frame at which the area under the ROC curve surpasses the value of 0.65 (upper blue line in panel B), and remains over this threshold for 15ms (vertical dashed lines in panel B; see materials and methods). The candidate SLR was identified if the target location was discriminated by the sEMG trace within the SLR epoch (grey patch) for both of the fast and slow trial sets. (C) Panel shows a line connecting the fast and slow DTs that are plotted for the slowest and fastest half of voluntary reaction times, and the line slope is showed. For this participant, both the early and late DTs are inside the SLR epoch (grey patch) and the line slope exceeds 67.5°, thus indicating the presence of a visuomotor response that is more time-locked to the stimulus onset than to the reaction time. (E) Panel shows the initiation time (dashed red line) obtained by running the ROC analysis on the full set of trials, and fitting a two-pieces “DogLeg” linear regression on the ROC curve to determine the point in time at which the ROC curve started to deviate positively toward the discrimination threshold (intersection point between the red lines; see materials and methods).

To quantify the SLR amplitude, on each trial we measured the mean sEMG activity recorded in the 10ms subsequent to the DT of the slow trial sets (Contemori et al. 2020). This method allowed us to quantify the muscle activity enclosed in a short time window in which the earliest target-related EMG response had been identified (i.e. DT within 80-120ms from target onset time) for both the fast and slow trial sets.

#### Cue-induced effect dimension

In this study, we expected to observe cue-induced modifications of the volitional and express visuomotor responses relative to control conditions. This would indicate that cue information was encoded by some neural circuit to bias the ensuing target-related response. We quantified the RT and SLR (initiation time and magnitude) differences between control and cue conditions both as absolute and percentage changes from control conditions and (termed as *cue-induced gain:* equation 1):

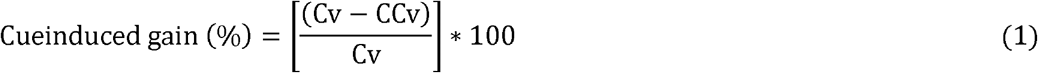

Where Cv represents the control value and CCv the cue condition value.

For the RT and SLR initiation time, we concluded that the cue exerted an advantaging effect if it led to shorter latencies than control (i.e. positive cue-induced gains). By contrast, we concluded that the cue exerted a disadvantaging effect if it led to longer latencies than control conditions (i.e. negative cue-induced gains). For the SLR magnitude, we inverted the order of members of the subtraction in equation 1: (*Cv* − *CCv*) → (*CCv* − *Cv*). This allowed us to index the cue-induced gain as positive (i.e. cue advantage effect) if the SLR size was larger in cue than control conditions, and negative (i.e. cue disadvantage effect) if the SLR had a larger magnitude in control than cue conditions.

#### Correlation of SLR magnitude with reaction time

One of the most intriguing questions about the putatively subcortical SLRs is whether or not they can contribute to volitional visuomotor behaviour. To disentangle the functional contribution of SLRs to voluntary movement initiation, we ran a correlation analysis between the SLR size and the corresponding RT on a trial-by-trial basis (Pruszynski et al. 2010; Gu et al. 2016; Contemori et al. 2020). The identification of a negative correlation between the SLR magnitude and RT across the different target conditions would indicate that the SLR size may influence the movement initiation, regardless of the type of stimulus (symbolic or low-contrast) cueing the target location.

#### Statistical analysis

Statistical analyses were performed in SPSS (IBMSPSS Statistics for Windows, version 25, SPSS Inc., Chicago, Ill., USA) and Matlab (version R2018b, TheMathWorks, Inc., 318 Natick, Massachusetts, United States). Results were analysed with t-test and repeated measure ANOVA models as the normality of the distributions was verified by the Shapiro-Wilk test. When ANOVA revealed a significant main effect or interaction, paired sample t-test were used for post-hoc comparisons. The chi-squared test was used to analyse changes in SLR prevalence between predicable and unpredictable conditions. For correlation analyses, the Pearson coefficient (r) was computed to index the strength of association between variables. For all tests, the statistical significance was designated at *p*< 0.05.

Formal within-participant statistical comparisons could not be conducted if SLRs occurred infrequently across the different target conditions. In this circumstance, we used a single-subject statistical analysis that aimed to test the reliability of the time-series ROC analysis to compare different stimulus conditions at the single-subject level (Contemori et al. 2020). Briefly, for each target condition we generated one thousand bootstrapped data sets from the original set of trials. We then ran the ROC and DogLeg analyses on each bootstrapped data set to extrapolate the distribution of SLR initiation time and magnitude. To test the statistical significance of the contrasts between the different target conditions, we compared one randomly re-sampled set of values from one target condition distribution with one randomly re-sampled set of values from the other target condition distribution (i.e. one thousand unique data comparisons for each of the three dependent variables). If the values for one target condition were larger or smaller than for the other target condition in more than 95% (i.e. >950) of cases, we concluded that the difference between the two target conditions was significant (for further details see supplementary materials in Contemori et al. 2020).

## RESULTS

### Experiment 1: symbolic cue

#### Task performance

A significant main effect of cue-condition on task correctness (*F_2,15_*=20.3, *p*<0.001) was obtained by running a one-way repeated measures ANOVA analysis. The post-hoc analysis (paired t-test) revealed that the prevalence of correct reaches was significantly lower in the invalid cue condition (78.3±16.1%) than the control (94.9±4.5%; t=4.6, *p*<0.001) and valid cue conditions (96.4±2.8%; t=4.5, *p*<0.001), whereas no significant difference was observed between the neutral and valid cue conditions. The fact that the highest error rate was observed with invalid cues suggests that the participants were biased to move toward the cued location. However, in the majority of invalid cue trials they correctly used the target spatial information to orient the final visuomotor response.

For the RT, we observed a significant main effect of cue-condition (one-way ANOVA: *F_2,15_*=27.6, *p*<0.001). The post-hoc analysis showed significantly shorter RTs for valid than control cue conditions (paired t-test: t=6.2, *p*<0.001; Figure 4A). By contrast, the RT was significantly longer with invalid than other cue conditions (paired t-test: control-invalid, t=3.3, *p*=0.003; valid-invalid, t=5.9, *p*<0.001; Figure 4A). Furthermore, validly cueing the target led to significantly positive percentage differences relative to control conditions (one sample t-test: t=6.4, *p*<0.001; Figure 4B), whereas significantly negative cue-induced percentage gains resulted from invalidly cueing the target (one sample t-test: t=3.1, *p*=0.004; Figure 4B). These findings indicate that the participants used the information extrapolated from the symbolic cue to improve their task performance.

**Figure 4:**
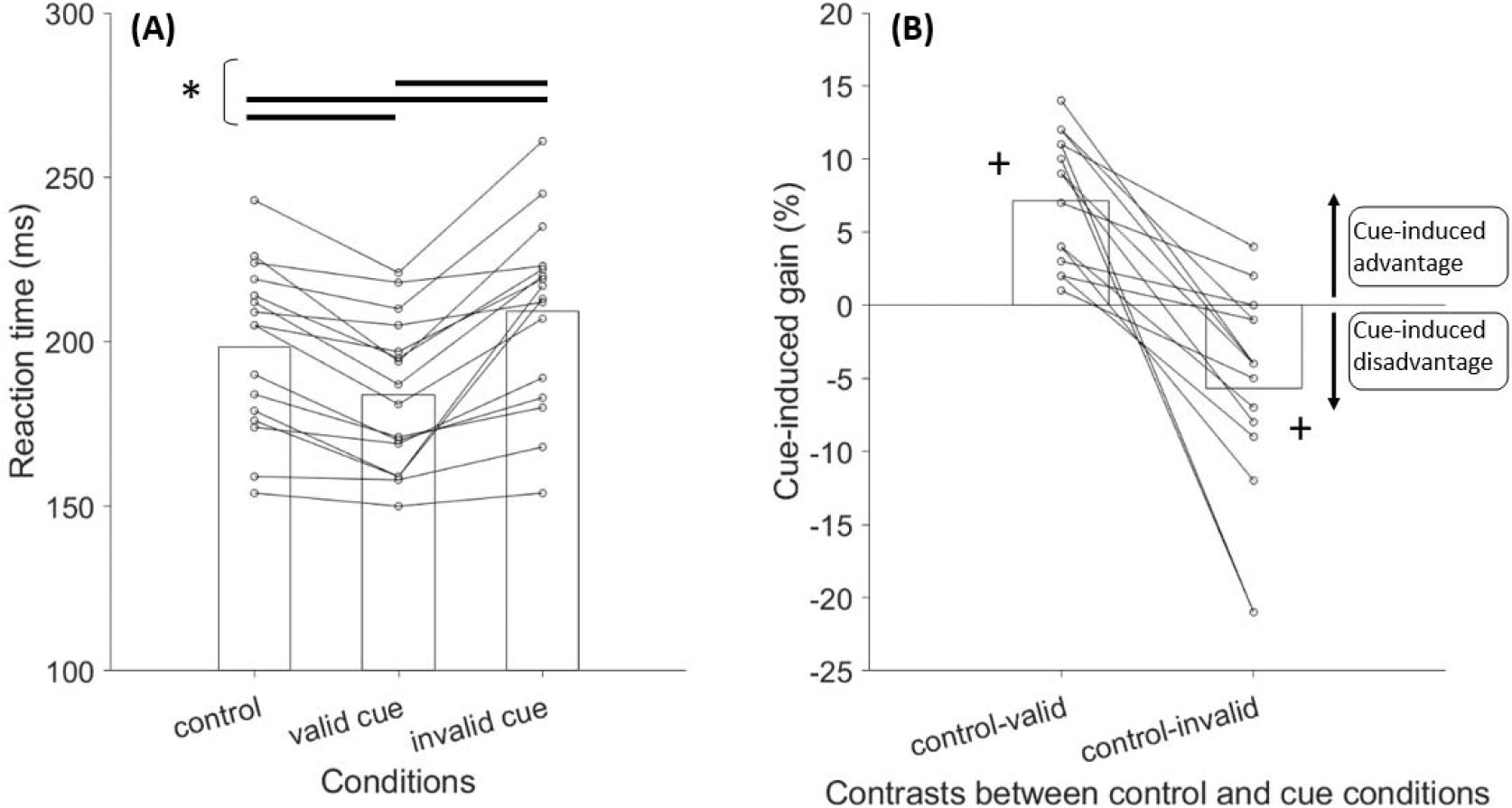
(A) Panel shows the latency of correct reaches in the control, valid and invalid cue conditions of the first experiment (see materials and methods). (B) Panel shows the percentage gains relative to control conditions induced by validly or invalidly cueing the target location with the arrow-shaped symbolic cues (see materials and methods). Positive cue-induced gains mean that cueing the target location advantaged the volitional movement initiation, whereas negative gains indicate disadvantaging cue-induced effects on reaction time. Each black line represents one participant, and the bars represent the mean values. Significant differences between task conditions: * *p*< 0.01. Significant difference from 0%: +*p*< 0.01.

#### Identified SLRs

To be classified as an SLR, the target location had to be discriminated from the sEMG signal within 80-120ms after the stimulus presentation in both fast and slow trial sets without, or with minimal, co-variation with the volitional RT (see materials and methods). For the PMch, the conditions for positive SLR detection were satisfied in both control and valid cue conditions in twelve out of sixteen participants, but only 6 of them also expressed an SLR in the invalid cue condition, and two participants did not express any SLR (Table 1). Notably, the valid cue condition promoted SLR generation among two participants who were otherwise negative SLR producers in the other task conditions (i.e. participants 3 and 13, Table 1). These observations resulted in significantly (*p*<0.05) lower SLR-prevalence for invalid cues than for control (chi-squared test; *p*=0.033, chi-squared=4.6, df= 1) and valid cue conditions (chi-squared test; *p*=0.003, chi-squared=8.5, df= 1). Notably, the high SLR prevalence in the control cue condition is consistent with recent studies (Kozac et al. 2020; Contemori et al. 2020) that used similar versions of the emerging target paradigm described here. This confirms the effectiveness of the paradigm for eliciting SLRs.

**Table 1:** Occurrences of positive SLRs (✓) in the clavicular head of the pectoralis major muscle (PMch) and the posterior deltoid (PD) across participants in all three cue conditions tested in experiment 1.

| Cue conditions | Control |  | Valid |  | Invalid |  |
| --- | --- | --- | --- | --- | --- | --- |
|  | PMch | PD | PMch | PD | PMch | PD |
| Participant |  |  |  |  |  |  |
| 1 | ✓ | ✓ | ✓ | ✓ | ✓ | - |
| 2 | ✓ | - | ✓ | - | ✓ | - |
| 3 | - | - | ✓ | - | - | - |
| 4 | ✓ | - | ✓ | - | - | - |
| 5 | ✓ | - | ✓ | ✓ | - | - |
| 6 | - | - | - | - | - | - |
| 7 | ✓ | - | ✓ | - | - | - |
| 8 | ✓ | - | ✓ | - | ✓ | - |
| 9 | ✓ | - | ✓ | - | - | - |
| 10 | ✓ | ✓ | ✓ | ✓ | ✓ | - |
| 11 | - | - | - | - | - | - |
| 12 | ✓ | - | ✓ | - | ✓ | - |
| 13 | - | - | ✓ | - | - | - |
| 14 | ✓ | - | ✓ | - | - | - |
| 15 | ✓ | - | ✓ | - | - | - |
| 16 | ✓ | - | ✓ | - | ✓ | - |
| Total SLRs (#) | 12 | 2 | 14 | 3 | 6 | 0 |
| SLR prevalence (%) | 75 | 12.5 | 87.5 | 18.75 | 37.5 | 0 |

The fact that many fewer SLRs were observed for the PD (Table 1) is consistent with the effects of isolated shoulder transverse extensor muscles preloading, which enhances the pre-target activity of the PMch but not that of the PD (Contemori et al. 2020). Given the low occurrence of SLRs for the PD, only the PMch was considered for statistical comparisons between the different cue conditions.

Cueing the target location influenced the timing and amplitude of SLRs. For the exemplar participant in figure 5, the sEMG signal started to deviate from baseline 87ms after target presentation for the valid cue condition, and at 95ms for the neutral cue condition (Figure 5C). For the invalid cue condition, the muscle started to encode the target location at 121ms from its presentation and, therefore, after the SLR epoch (Figure 5C). Furthermore, SLR magnitude was larger for the valid (76μV) than neutral (55μV) cue conditions. These findings resulted in positive cue-induced SLR initiation time (8.4%) and magnitude (38.2%) gains, relative to control conditions.

**Figure 5:**
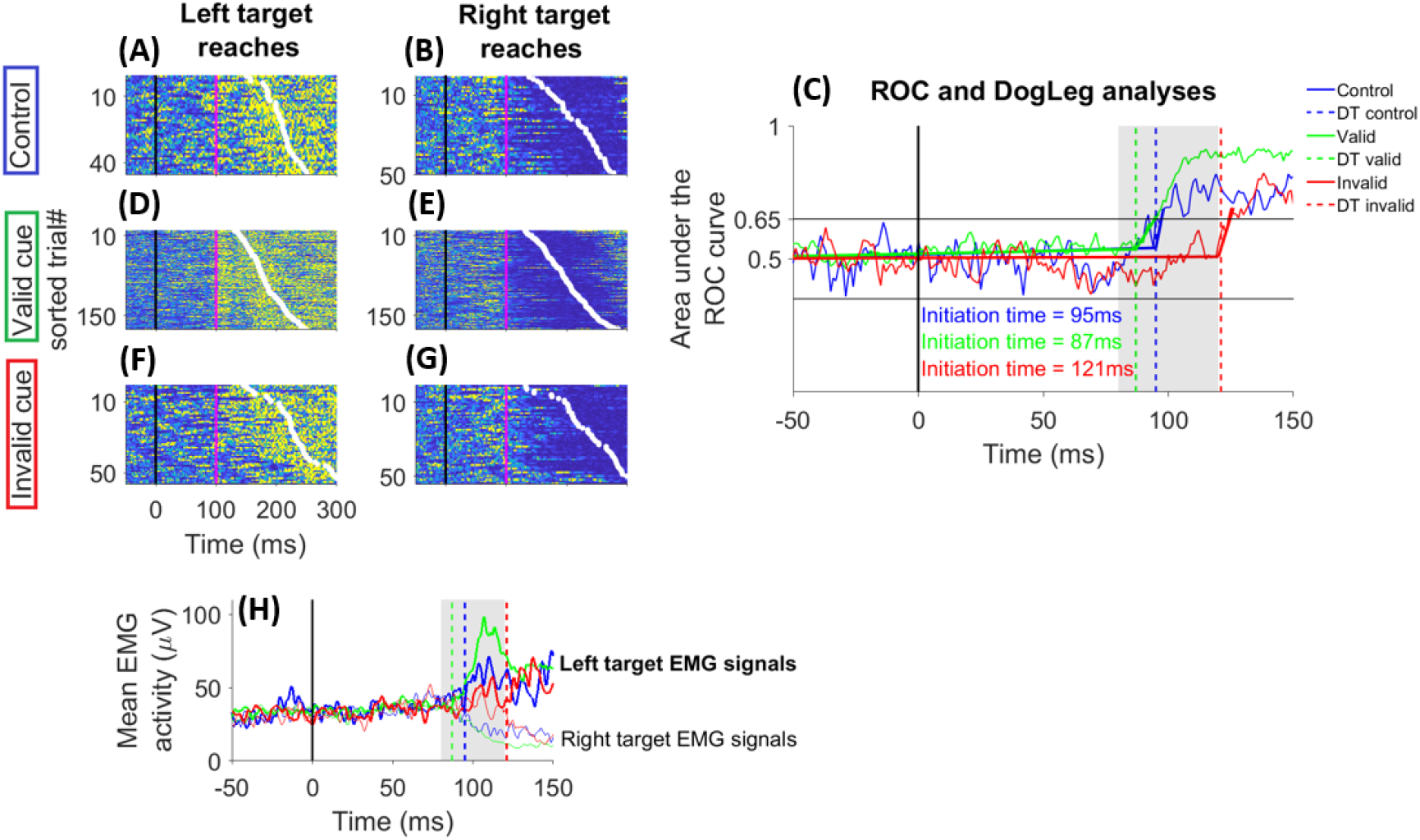
Surface EMG activity of the pectoralis major clavicular head muscle of an exemplar participant who completed the first experiment, and exhibited an SLR in control and valid cue conditions, but not in invalid cue conditions (participant 5, Table 1). For each cue condition, rasters of rectified sEMG activity from individual trials are shown (A, B, D-G; same format as figure 2). The solid magenta line indicates the expected initiation time of the SLR (~100ms from target onset). (H) Panel offers a zoomed view of the mean sEMG activity (thick lines = left target reaches; thin lines = right target reaches), and the vertical dashed lines show the initiation time of the target-related muscle response. The initiation time was indexed as the point in time at which the ROC curve started to positively diverge toward the 0.65 discrimination threshold (see materials and methods). Panel C offers a zoomed view of ROC and DogLeg analyses that were run to index the initiation time of the target-related EMG response. For this participant, the ROC curve starts to deviate earlier in valid (87ms, intersection between the straight green lines) than control (95ms, intersection between the straight blue lines) cue conditions, and after the SLR epoch in invalid cue conditions (121ms, intersection between the straight red lines).

Similar trends were observed across the 12 participants who produced an SLR to the control and valid cue conditions (Table 1). The initiation time was significantly shorter, and the SLR magnitude significantly larger, in the valid (~85±8ms, ~66±32μV) than control (~95±10ms, ~59±33μV) cue conditions (paired t-test: initiation time, t=4.1, *p*<0.001; magnitude, t=1.8, *p*=0.003; figure 6A and C). In addition, we observed significantly positive cue-induced percentage gains for each of the SLR parameters (one sample t-test: initiation time, t=4.6, *p*<0.001; magnitude, t=2.1, *p*=0.001), relative to the control condition (inset plots in figure 6A and C). These results indicate a cue-induced SLR facilitation relative to control conditions when the target appeared at the expected location.

**Figure 6:**
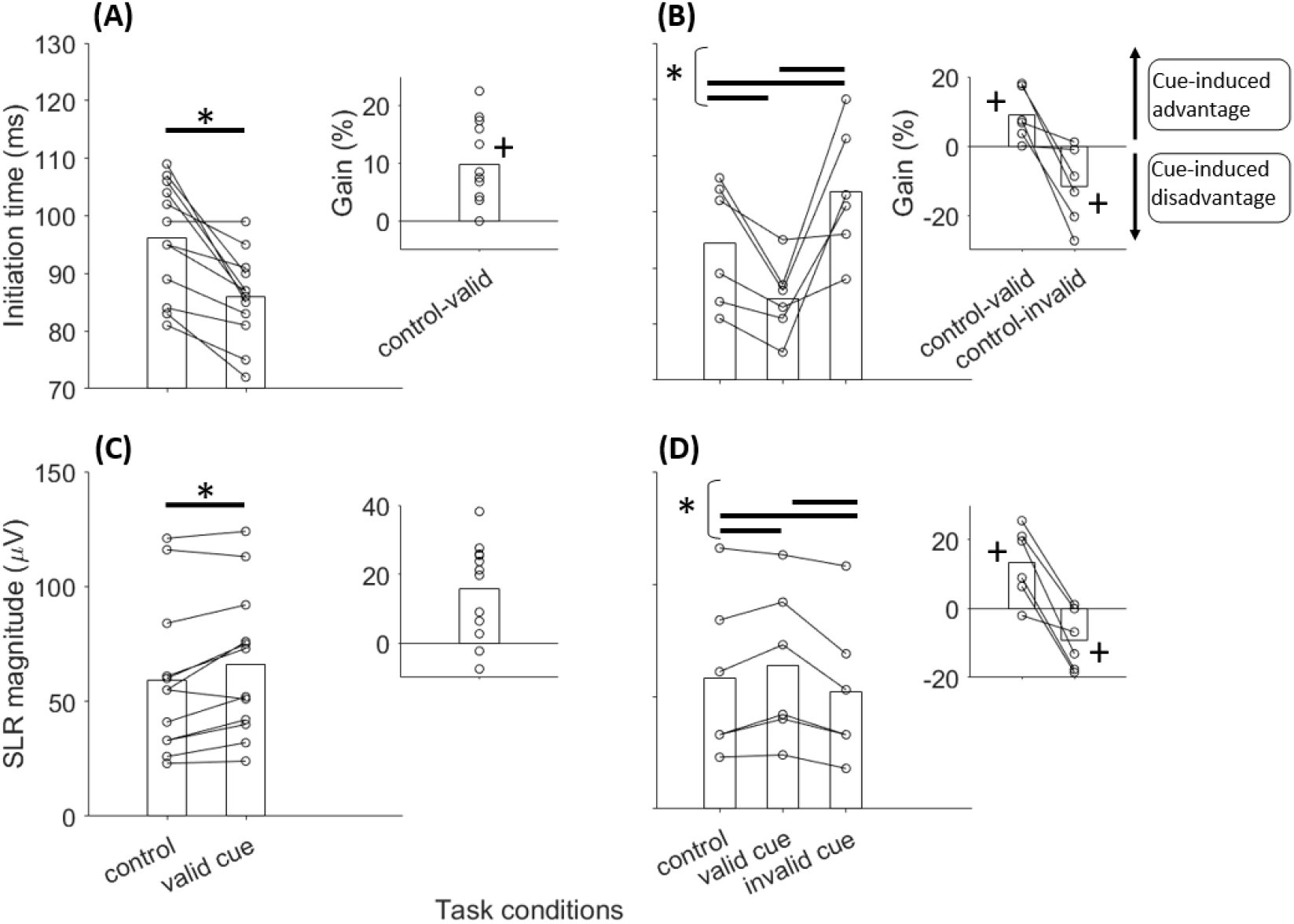
Latencies and magnitude of the express visuomotor responses in the first experiment. Panels A and C show the results from twelve participants who exhibited an SLR in control and valid cue conditions (see Table 1), and the inset panels show the percentage gain induced by validly cueing the target location relative to control conditions. Panels B and D show the results of six participants who exhibited an SLR in control, valid and invalid cue conditions (see Table 1), and the inset panels show the percentage gain induced by validly and invalidly cueing the target location relative to control conditions. Positive cue-induced gains mean that cueing the target location advantaged the SLR expression, whereas negative gains mean disadvantaging cue-induced effects. Each solid black line and dot represent one participant, and the bars represent the average across participants. Validly cueing the target location with the symbolic arrow cue led to significantly (*p<0.01) faster (A) and larger (C) SLRs than control conditions, and to significantly positive (+p<0.01) percentage gains relative to control conditions (inset plots in A and C panels). The second column shows that the SLRs were significantly (*p<0.05) faster (B) and stronger (D) than control with valid cues, and significantly (*p<0.05) slower (B) and smaller (D) than control with invalid cues. Moreover, validly cueing the target location led to significantly (+p<0.05) positive percentage gains relative to control conditions, whereas significantly (+p<0.05)

To complete the description of cue-induced effects on SLR expression, we ran a one-way repeated measure ANOVA analysis on the 6 participants who exhibited an SLR among all three cue conditions (Table 1). For this analysis, we defined the cue-validity (3 levels: neutral, valid, invalid) as within-participant factor. A significant cue-validity main effect was found for initiation time (*F_2,5_*=10.3, *p*=0.004) and SLR magnitude (*F_2,5_*=9.87, *p*=0.004). Post-hoc analyses showed significantly longer SLR initiation times with invalid than other cue conditions (paired t-test: control-invalid, t=2.8, *p*=0.019; valid-invalid, t=3.5, *p*=0.008; figure 6B and D). The SLR size was significantly smaller with invalid than other cue conditions (paired t-test: control-invalid, t=2.4, *p*=0.03; valid-invalid, t=3.6, *p*=0.008). The results for the percentage change from control were consistent with the absolute comparisons. More precisely, we observed significantly negative cue-induced gains with the invalid relative to control cue conditions (one sample t-test: initiation time, t=2.6, *p*=0.025; magnitude, t=2.6, *p*=0.024; inset panels in figure 6B and D). These results suggest SLR inhibition effects when the expected and actual target locations were mismatched.

### Experiment 2: low-contrast cue

#### Task performance

The occurrence of correct reaches was ~95% for control and valid low-contrast cue conditions, ~90% in the invalid low-contrast cue condition and ~85% for the single low-contrast target condition. The one-way repeated measures ANOVA analysis showed a main effect for task condition (*F_2,11_*= 4.9, *p*=0.007). The post-hoc analysis evidenced a significantly lower correct response rate for the low-contrast target than the control (paired t-test: t=4.3, *p*=0.001) and valid cue (paired t-test: t=-3.7, *p*=0.003) conditions, whereas no significant difference was observed between the invalid cue and other task conditions. These results suggest that target detection was impaired, but not fully obliterated, by the presentation of stimuli that were around the threshold for correct detection. Furthermore, the data indicate that participants moved correctly toward the high-contrast target even when it was preceded by the low-contrast cue at the opposite location.

A significant task-condition main effect (one-way ANOVA: *F_2,15_*= 27.6, *p*<0.001) was found for RT. The RT was significantly longer in the low-contrast than in all of the other target conditions (paired t-test: control-low contrast, t=5.9, *p*<0.001; low contrast-valid, t=6.4, *p*<0.001; low contrast-invalid, t=4.3, *p*<0.001; Figure 7A). Further, the RT was significantly longer for the invalid cue condition than the control (paired t-test: t=3.1, *p*=0.005) and valid cue conditions (paired t-test: t=4.7, *p*<0.001). Finally, validly cueing the target led to significantly faster RTs than control conditions (paired t-test: t=5.4, *p*<0.001; Figure 7A). The absolute cue-induced changes were consistent with the percentage cue-induced gains relative to control conditions. More precisely, the valid cue led to significantly positive RT gains relative to control (one sample t-test: t=6.2, *p*<0.001) conditions, whereas significantly negative RT gains were observed with invalid cues (one sample t-test: t=3.2, *p*=0.004; Figure 7B). These findings indicate that the low-contrast stimulus biased the volitional reaching behaviour despite its low saliency for movement initiation, its temporal proximity (~24ms) to the high-contrast target and its lack of predictive value (50% validity) for signalling the location of the high-contrast target.

**Figure 7:**
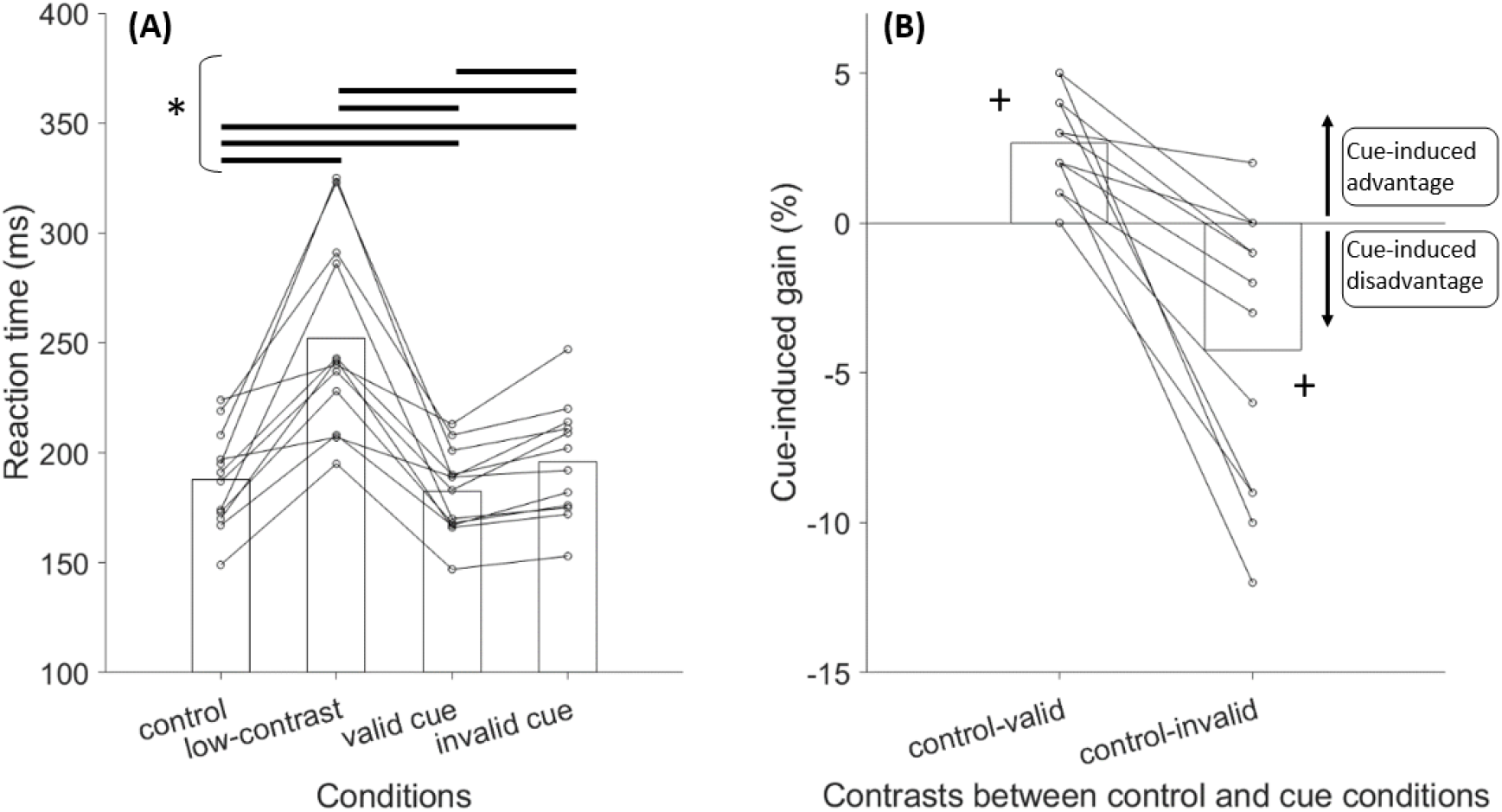
(A) Latency of correct reaches toward high-contrast targets (control condition), low-contrast targets, and high-contrast targets cued by low-contrast stimuli appearing at the same (valid cue) or opposite (invalid cue) location. (B) Panels shows the percentage gains relative to control conditions induced by validly or invalidly cueing the target location with the low-contrast cues (same format as figure 4). Significant differences between task conditions: * *p*< 0.01. Significant difference from 0%: +*p*< 0.01.

#### SLRs

The second experiment was completed by 12 participants who also participated in the first experiment. In ten of them, we detected an SLR on the PMch muscle either when the high-contrast target appeared alone (control condition) or when it was validly cued by the low-contrast stimulus, but only five of them had an SLR also for the invalid cue condition (Table 2). The presentation of the low-contrast stimulus alone elicited an SLR in only two participants, who also had an SLR in the control and valid cue conditions, but not in the invalid cue condition (see participants 1 and 3 in Table 2). Finally, two participants did not exhibit any SLR (i.e. participants 4 and 8, Table 2). Akin to the first experiment, a sufficient number of SLRs for statistical comparisons between the target conditions was obtained only for the PMch muscle (Table 2).

**Table 2:** Occurrences of positive SLRs (✓) in the clavicular head of the pectoralis major muscle (PMch) and the posterior deltoid (PD) across participants in all four task conditions tested in experiment 2. Participants 1-12 correspond to participant 8, 7, 1, 11, 9, 4, 10, 15, 12, 14, 5 and 13 in table 1.

| Task conditions |  | Control |  | Low-contrast |  | Valid |  | Invalid |  |
| --- | --- | --- | --- | --- | --- | --- | --- | --- | --- |
| Muscles |  | PMch | PD | PMch | PD | PMch | PD | PMch | PD |
| Participant |  |  |  |  |  |  |  |  |  |
| 1 |  | ✓ | - | ✓ | - | ✓ | - | - | - |
| 2 |  | ✓ | - | - | - | ✓ | - | ✓ | - |
| 3 |  | ✓ | - | ✓ | - | ✓ | - | - | - |
| 4 |  | - | - | - | - | - | - | - | - |
| 5 |  | ✓ | - | - | - | ✓ | - | ✓ | - |
| 6 |  | ✓ | - | - | - | ✓ | - | - | - |
| 7 |  | ✓ | ✓ | - | - | ✓ | ✓ | ✓ | - |
| 8 |  | - | - | - | - | - | - | - | - |
| 9 |  | ✓ | - | - | - | ✓ | - | - | - |
| 10 |  | ✓ | - | - | - | ✓ | - | - | - |
| 11 |  | ✓ | - | - | - | ✓ | - | ✓ | - |
| 12 |  | ✓ | - | - | - | ✓ | - | ✓ | - |
| Total SLRs (#) |  | 10 | 1 | 2 | 0 | 10 | 1 | 5 | 0 |
| SLR prevalence (%) |  | 83.3 | 8.3 | 16.7 | 0 | 83.3 | 8.3 | 41.7 | 0 |

Given that the same ten participants expressed an SLR to control and valid cue conditions (i.e. participants 1-3, 5-7 and 9-12, Table 2), we only considered the control condition to test whether the SLR prevalence was significantly different across conditions. The Chi-squared test returned a significantly higher (*p*<0.05) SLR prevalence for control than both low-contrast target (*p*=0.001, chi-squared=10.7, df= 1) and invalid cue conditions (*p*=0.035, chi-squared=4.4, df= 1). This suggests that the low-contrast target was a less salient stimulus for SLR generation than the high-contrast target. Further, cueing the high-contrast target with an invalid low-contrast cue impaired, but did not completely obliterate, the SLR expression.

Figure 8 shows the results of one exemplar participant who participated in the second experiment (i.e. participant 12, Table 2). For this participant, the ROC curve started to deviate from chance earlier for the valid (81ms; Figure 8I) and later for the invalid (110ms; Figure 8L) cue relative to control conditions (97ms; Figure 8C). By contrast, in the low-contrast target condition the sEMG signal started to encode the location in 130ms after the stimulus presentation (Figure 8F), thus after the SLR epoch (i.e. 80-120ms after stimulus onset time). The size of the SLR was similar between the high-contrast target (28μV) and valid cue conditions (25μV), whereas a smaller SLR magnitude was observed for the invalid cue condition (16μV).

**Figure 8:**
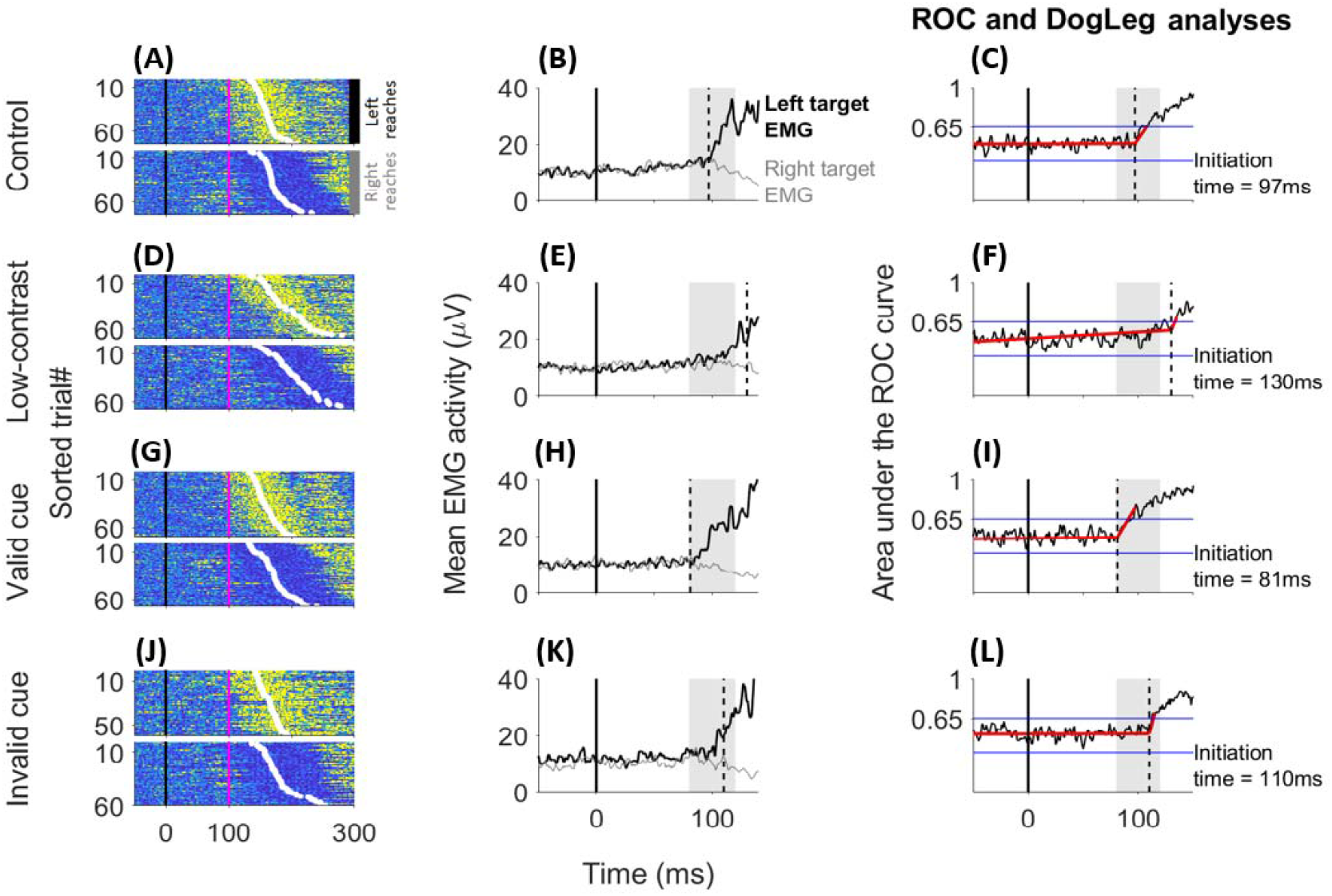
Surface EMG activity of the pectoralis major clavicular head muscle of an exemplar participant who completed the second experiment, and exhibited an SLR in (A) control, (G) valid and (J) invalid cue conditions, but not in (D) low-contrast target condition (participant 12, Table 2). For each condition, rasters of rectified sEMG activity from individual trials are shown (A, D, G, J; same format as figure 5). Panels B, E, H and K offer a zoomed view of the mean sEMG activity, and the vertical dashed lines show the initiation time of the target-related muscle response (see materials and methods; same format as figure 5).For this participant, the ROC curve starts to deviate at 97ms in (C) control, 81ms in (I) valid and 110ms in (L) invalid cue conditions, whereas the initiation time in (F) low-contrast target condition was at 130ms and, thereby after the SLR epoch (grey patch).

Similar trends were observed across the 10 participants who expressed an SLR in control and valid cue conditions (Table 2). More precisely, the SLR initiation time was significantly earlier for the valid (~81±2ms) cue than control (~90±5ms) conditions (paired t-test: t=6.1, *p*<0.001; Figure 9A). Furthermore, we observed a significantly positive cue-induced percentage gain of the initiation time relative to the control condition (one sample t-test: t=6.7, *p*<0.001; inset plot in Figure 9A). By contrast, no significant difference was found between the valid cue and control conditions for the SLR magnitude (Figure 9C). These results suggest that the SLR latency can be shortened by the presentation of a low-contrast stimulus appearing shortly in advance of, and at the same location, as a high-contrast target.

**Figure 9:**
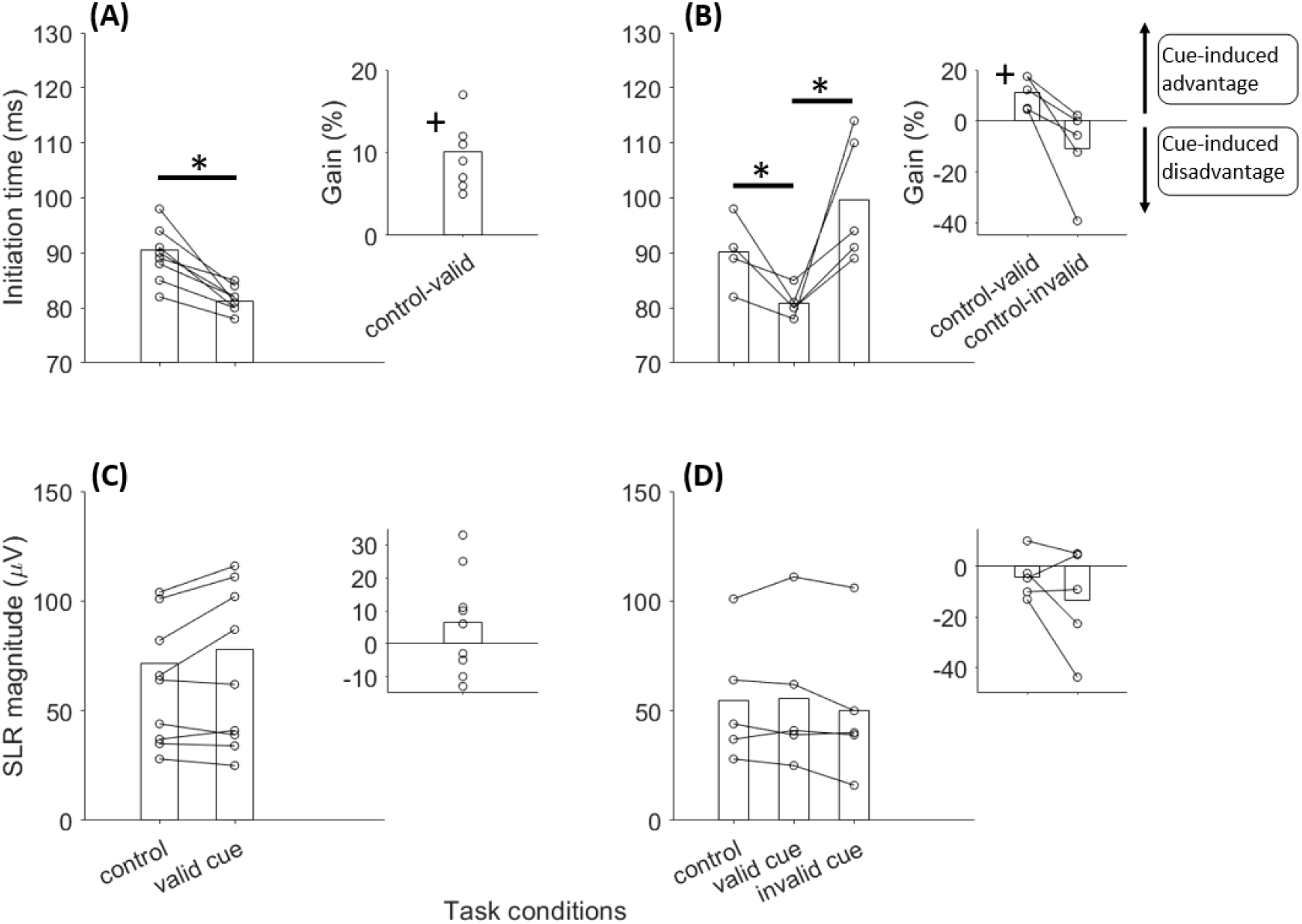
Latencies and magnitude of the express visuomotor responses in the second experiment. The first column of panels shows the results often participants who exhibited an SLR in control and valid cue conditions (see Table 2). The second column of panels shows the results of five participants who exhibited an SLR in control, valid and invalid cue conditions (see Table 2).Validly cueing the target location with the low-contrast cue led to significantly faster SLRs than control condition (A, * *p*<0.01; B, * *p*<0.05), and to a significantly positive cue-induced percentage gain relative to control condition (inset plot in A, + *p*<0.01; inset plot in B, + *p*<0.05). Further, valid low-contrast cues led to significantly (* *p*<0.05) faster SLRs than invalid cue conditions (B).

The exemplar participant’s results (Figure 8) were also consistent across the five participants who exhibited an SLR in the high-contrast, valid cue and invalid cue conditions (i.e. participants 2, 5, 7, 11 and 12, Table 2). For these participants, we ran a one-way ANOVA analysis with task-condition (3 levels: control, valid cue, invalid cue) as within-participant factor. A significant task-condition main effect was found for the initiation time (*F_2,4_*=6.9, *p*=0.018), but not for the SLR magnitude (*p*=0.213). Post-hoc analysis showed significantly faster SLRs with the valid than invalid cue conditions (paired t-test: t=3.3, *p*=0.015; Figure 9B). The SLR latency was also ~10ms shorter in control than invalid cue conditions (Figure 9B), but this difference was not statistically significant (paired t-test: t=1.5, *p*=0.1). Invalid low-contrast cues led to negative percentage gains of SLR timing (~ −11%, inset plot in Figure 9B) and magnitude (~ −13%, inset plot in Figure 9D) relative to control conditions. However, the one-sample t-test did not show significant contrasts (initiation time, t=1.5, *p*=0.11; SLR magnitude, t=1.4, *p*=0.11), probably because of the small sample size. These findings suggest that cueing the location of high-contrast targets with barely detectable cues can modulate the SLR expression as a function of the compatibility between the two stimuli positions.

In figure 10 are shown the data of one participant (S1) who produced an SLR in control, low-contrast target and valid cue conditions, but not in the invalid cue condition (i.e. participant 1, Table 2). A similar SLR distribution was observed in only one other participant (S2) of the second experiment (i.e. participant 3, Table 2). Given that only two participants exhibited an SLR for the low-contrast target condition, we ran a single participant statistical-analysis to test the significance of the contrasts between the target conditions (see materials and methods; Contemori et al. 2020). Participant S1 had a median initiation time of 97ms and a 95% confidence interval of [90-104] for control target, 112ms [102-122] for low-contrast target and 81ms [73-90] for valid cue conditions. The SLR magnitude was 42μV [38-46] for control target, 28μV [21-35] for low-contrast target and 41μV [36-46] for valid cue conditions. For participant S2, the initiation time was 94ms [88-100] for control target, 112ms [104-120] for low-contrast target and 84ms [77-91] for valid cue conditions. The SLR magnitude was 78μV [54-102] for control target, 48μV [24-72] for low-contrast target and 85μV [72-92] for valid cue conditions. For both participants, the initiation time was significantly shorter (*p*<0.05) with the valid cue condition than both control and low-contrast target conditions, and significantly longer than control with the low-contrast target condition. The SLR magnitude was significantly larger (*p*<0.05) with the valid cue than low-contrast target conditions. The size of the SLR was also larger in the control than low-contrast target conditions, but this difference was statistically significant (*p*<0.05) only for S1 (i.e. participant 1, Table 2). By contrast, for both participants the SLR size was not significantly different (*p*>0.05) between the control and valid cue conditions. These results indicate that some participants are capable of producing SLRs both to high-contrast and low-contrast stimuli. However, low-contrast targets have less saliency for the generation of rapid and large SLRs as compared with high-contrast targets. Further, the data confirm the advantaging effects of valid and low-contrast cues and, conversely, the negative effects of invalid low-contrast cues relative to control conditions.

**Figure 10:**
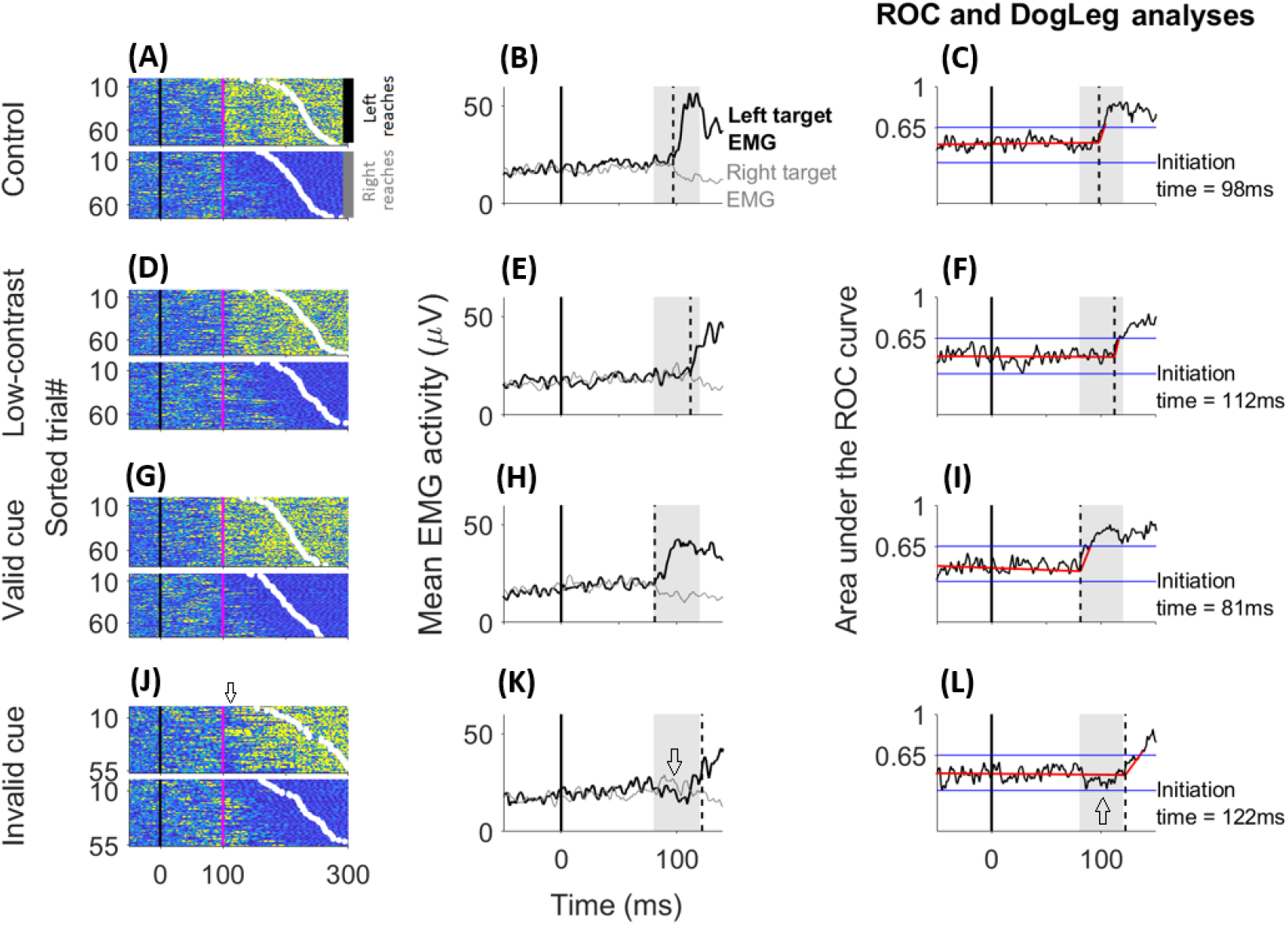
Surface EMG activity of the pectoralis major clavicular head muscle of a participant who exhibited an SLR in (A) control, (D) low-contrast target and(G) valid conditions, but not in (J) invalid cue condition (participant 1, Table 2). For each condition, rasters of rectified sEMG activity from individual trials (panel A, D, G and J), mean EMG traces (panel B, E, H and K) are shown, as are the outcomes of the time-series ROC and DogLeg linear regression analyses (panel C, F, I and L; same format as figure 8). For this participant, the ROC curve starts to deviate at 98ms in (C) control, 112ms in (F) low-contrast target and 81ms in (I) valid cue conditions, whereas the initiation time in (L) invalid target condition is at 122ms and, thereby after the SLR epoch (grey patch). In panel J, the arrow indicates short latency responses at ~100ms that are consistent with the low co-contrast cue location, before the muscle started responding to the high-contrast target. These rapid responses reflect the short-latency (~100ms) EMG activation for right targets and inhibition for left targets of the average EMG signal (arrow inside the grey patch in panel K), and underlies the negative deflection below 0.5 chance level of the ROC curve within the SLR epoch (arrow inside the grey patch in panel L).

In figure 10J, short-latency responses can be observed at ~100ms in the invalid cue trials before the muscle started responding to the high-contrast target (arrow in figure 10J). This reflects the erroneous activation/inhibition of the PMch and underlies the negative deflection below 0.5 chance level of the ROC curve within the SLR epoch (arrow inside the grey patch in figure 10K and L). Some express motor signals encoding the low-contrast cue location appear to have been delivered to the muscles. Such express visuomotor responses to a barely detectable stimulus might then be rapidly overridden by a response to a more salient target, at least when both visual events occur within a short temporal interval. This hypothesis remains tentative, however, because this phenomenon was observed in only one participant.

### Correlation analyses

#### Correlating reaction time with SLR magnitude

To disentangle the SLR contribution to volitional reaching behaviour, we tested the correlation between SLR magnitudes and RTs. Figure 11A shows this correlation for an exemplar participant (i.e. participant 2, Table 2). A negative RT x SLR magnitude correlation was found consistently among the SLR observations in the first (one sample t-test; control, t = 10.5, *p*<0.001; valid cue, t = 7.3, *p*<0.001; invalid cue, t = 6.9, *p*<0.001;Figure 11B)and second experiments (one sample t-test; control, t =8.5, *p*<0.001; valid cue, t =7.2, *p*<0.001; invalid cue, t =7.7, *p*<0.001; Figure 11C). A significant negative correlation was also observed for the two participants (S1, participant 1, Table 2; S2, participant 3, Table 2) who exhibited an SLR to the low-contrast targets (Pearson correlation coefficient (*r*): S1,*r* = −0.27, *p*=0.009; S3, *r* = −0.49, *p*<0.001).These findings are consistent with previous work (Pruszynski et al. 2010; Gu et al. 2016; Contemori et al. 2020), suggesting that the SLR contributes functionally to the volitional initiation of target-directed reaches regardless of how each is modulated by cues.

**Figure 11:**
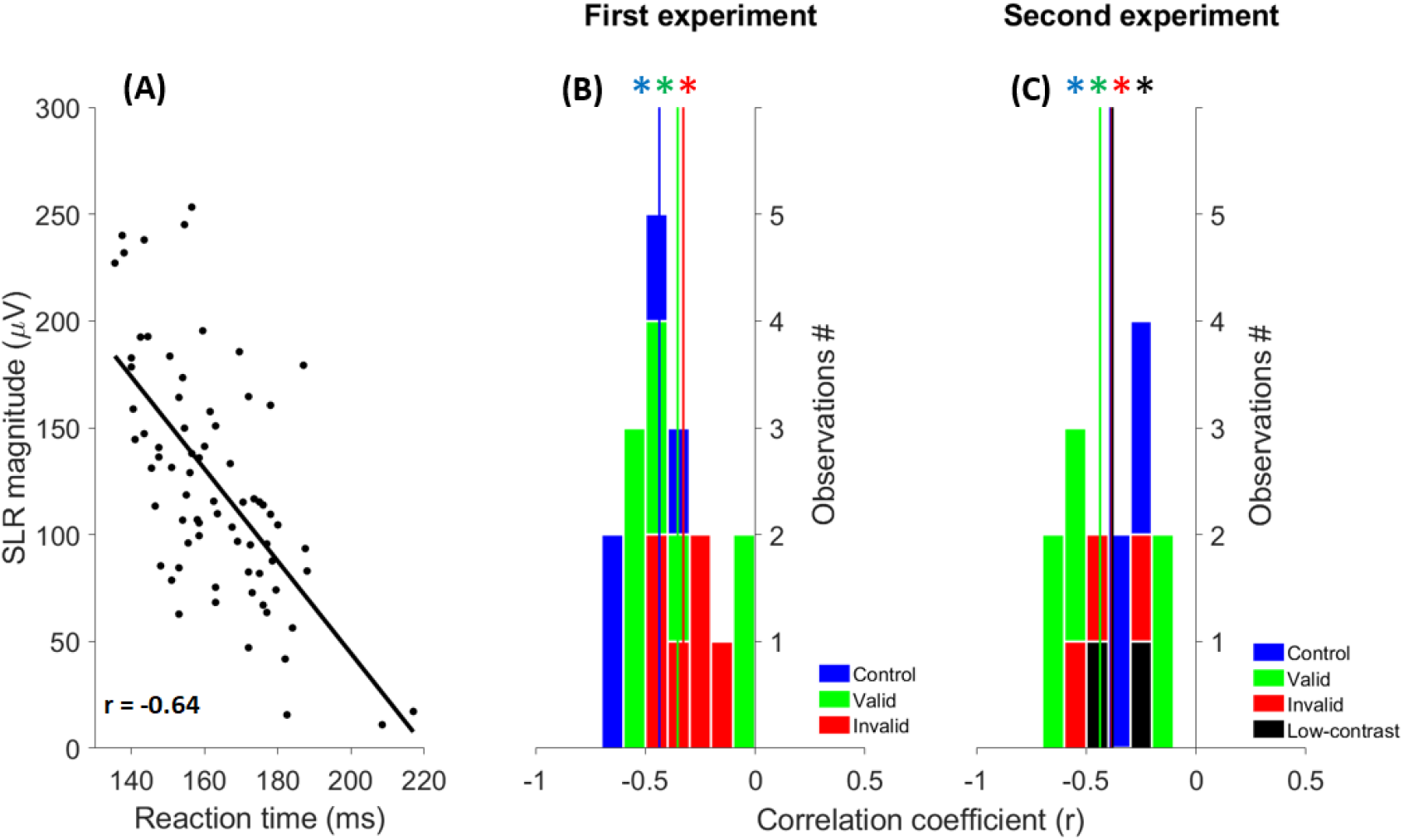
(A) Correlation between the reaction time and SLR magnitude from the pectoralis major clavicular head for an exemplar participant who expressed an SLR in the second experiment valid cue condition (participant 2, Table 2). Each data point represents a single trial and the solid blackline is the linear regression function. (B) Group correlation coefficient for all participants with at least an SLR in control (12 participants), valid (14 participants) or invalid (6 participants) cue conditions of the first experiment (see Table 1). (C) Group correlation coefficient for all participants with at least an SLR in control (10 participants), valid cue (10 participants), invalid cue (5 participants) or low-contrast target (2 participants) conditions of the second demonstrates a significant negative correlation (* p<0.01) with the movement initiation, irrespective of cueing the target location with symbolic (first experiment, B) or low-contrast cues (second experiment, C).

## DISCUSSION

### Experiment 1: Symbolic cue

In this study, the reaching task required rapid identification of the target location relative to hand position in order to program the reaching direction and associated coordination between the agonist\antagonist muscles. The arrow-shaped cue provided symbolic, but not spatial, information regarding the future target location because its position was irrelevant with respect to the two possible target locations. That is, the target position could be predicted only via a cognitive extrapolation of the arrow orientation. When this information was valid, the RT was shorter than in control conditions. However, this cue-induced benefit turned into a behavioural cost (i.e. delaying RT) when the cue was invalid. These observations are consistent with an overt *attention orientation* mechanisms (Posner 2016) that reflects cortical perception about the expected task.

In mammalian species, the neural networks involved in cortical attention orientation comprise complex feedback loops between prefrontal, parietal and sensory cortices and thalamic, basal ganglia and brainstem structures (for review see Baluch and Itti 2011; Knudsen 2018). For instance, Moore and Armstrong (2003) showed that microstimulation of the frontal eye field (FEF) enhanced neural activity of V4 area in monkeys. Further, the enhanced activity in V4 area was restricted to visual neurons encoding the visual field corresponding to the saccade that could be triggered by the FEF neurons undergoing the stimulation procedure. This suggests a cortico-cortical modulation mechanism by which higher-level premotor and motor areas can modify the activity of sensory cortices, such as those deputed to the processing of visual information. The symbolic cue-induced RT advantages may underlie priming mechanisms of the visual neurons encoding the cued location, consistent with an endogenous prioritization to sensory events occurring at the expected location. By contrast, the neural populations encoding the non-cued locations could be disengaged by suppressing cortico-cortical feedback signals (Baluch and Itti 2011; Knudsen 2018). This may result in a longer time to override the cue-driven expectation and transform the unexpected stimulus in the corresponding target-directed reach, consistent with the increase of volitional RTs with the invalid symbolic cues.

The prior information extrapolated from the symbolic cue also influenced the temporal and magnitude components of the SLR. Specifically, validly cueing the target location reduced the SLR initiation time and enlarged the SLR amplitude as compared to control conditions, whereas the opposite was observed with invalid symbolic cues. The SLR is the biomarker of a neural network that can rapidly generate muscle responses, which are computed in a hand-centric reference frame (Gu et al. 2018). This neural network may include the midbrain superior colliculus and its downstream connections with the brainstem reticular formation, which then projects to interneurons and motoneurons in the spinal cord. It is noteworthy that the existence of a subcortical network operating rapid visuomotor transformations in humans would indicate that the sensorimotor transformation of visual events is not an exclusive duty of high-level cortical sensorimotor areas. Given that the symbolic cue required cognitive extrapolation, we propose that the cue-induced SLR modifications reflect a cortical top-down modulation of the putative subcortical SLR network, including the superior colliculus.

The superior colliculus contribution to SLR generation is supported by evidence of collicular involvement in the production of express saccades (Dorris et al. 1997). This midbrain structure receives direct retinal inputs, but is also mutually interconnected with cortical areas responsible for the cascade of neural operations that transforms visual events into motor actions (i.e. visual, parietal and frontal cortices; Boehnke and Munoz, 2008). Peel et al. (2017) reported activity decrements of the superior colliculus neurons when the frontal-eye-field in monkeys was cryogenically inactivated. More recently, Dash et al. (2018) showed that FEF inactivation correlated with reduced occurrence of express saccades relative to control conditions. Critically, these findings indicate that the cortical top-down signals to the superior colliculus can modulate the express visuomotor transformations operated by this midbrain structure.

Cortical signals encoding cognitive expectations can be conveyed to the neural structures responsible for low-level processing and the rapid sensorimotor transformation of visual inputs, such as the superior colliculus. Selectively manipulating the activity of the topographically organized collicular visual map according to expected locations may increase the response to congruent sensory events and diminish the response to unexpected stimuli. For example, preceding work has shown that the presentation of temporally and spatially predictable targets facilitated the initiation of target-directed saccades within the express range (~100ms; Paré and Munoz 1996; Dorris et al. 2007). This suggests a contribution of cognitive expectation to the generation of express visuomotor responses. Moreover, expecting a stimulus to occur at a defined position correlates with inhibition of activity of the superior colliculus neurons encoding the locations distant from the saccadic goal (Dorris and Munoz 1998). This suggests that rapid collicular visuomotor transformations are modulated as a function of the pre-target collicular activity, which can be biased by cortical top-down signals originating from expectations about future sensory events. This cortical top-down priming might underlie a top-down attention orienting mechanism to increase the saliency of expected stimuli on the collicular visual map and to inhibit the responses to unexpected targets (Baluch and Itti 2011). Noteworthy, the cortical top-down SLR modulation hypothesis is consistent with recent evidence of SLR facilitation induced by temporal stimulus predictability and by briefly flashed stimuli, which activate both ON and OFF responses in superior colliculus (Contemori et al. 2020). This neural mechanism may underlie the faster and larger SLRs observed when the target appeared in an expected location, and the slower and smaller SLRs expressed with invalid cues relative to control conditions.

### Experiment 2: Low-contrast cue

The low-contrast targets had a low saliency for both volitional and express visuomotor behaviours, which underlies both the delayed RT and impaired SLR expression relative to control conditions. Only two participants exhibited an SLR for the low-contrast target condition (participants 1 and 3, Table 2) and it was delayed and smaller than that expressed with the high-contrast target condition. These results are consistent with previous work showing that both visual responses in the superior colliculus (Marino et al. 2010) and the SLR (Wood et al. 2015) are delayed as the target-to-background contrast is reduced.

Despite its low saliency, the low-contrast stimulus led both to volitional and express behaviour modulations when it was used as a cue for the high-contrast target. Specifically, the valid low-contrast cues reduced both the RT and SLR latency relative to control conditions, whereas the invalid cues led to the opposite effects. Further, invalid low-contrast cues obliterated the SLR in five out of ten participants who exhibited it in control and valid cue conditions (Table 2). These phenomena are unlikely to originate from the same neural mechanisms proposed for the symbolic cue effects. The symbolic cue was predictive for target location (i.e. 75% validity) and required cortical extrapolation of the arrow orientation, which we enabled experimentally by a CTOA >1s. By contrast, the low-contrast cues were designed to minimize cortical involvement by their low saliency, brief CTOA (~24ms) and irrelevant validity (50%). This is consistent with the low (~10%) occurrence of incorrect (i.e. cue-directed) reaches in the invalid cue conditions, which indicates that participants moved toward the high-contrast target even when it was invalidly cued by the low-contrast cue appearing in the opposite visual hemi field. Therefore, the SLR consequences of barely detectable cues likely originated from neural circuits operating low-level visual processing and visuomotor transformations, rather than cortical visuomotor networks.

The superior colliculus is known to perform low-level processing and short-latency visuomotor transformation of visual events detected by the retinal photoreceptors (Boehnke and Munoz 2008; Gandhi and Katnani 2011; Basso and May 2017). Furthermore, this midbrain structure is proposed to contribute to mechanisms of bottom-up attention orientation (Baluch and Itti 2011; Knudsen 2018). The bottom-up attention evolves rapidly after a sensory event and is exclusively sensitive to the physical attributes of the stimulus, such as its spatial location (Baluch and Itti 2011). Neural correlates of bottom-up attention orientation in the superior colliculus have been reported in non-human primates, and there is some evidence that perturbations of superior colliculus activity can influence both conscious perception and volitional motor behaviour (Baluch and Itti 2011; Corneil and Munoz 2014; Knudsen 2018). For instance, Muller et al. (2005) showed that microstimulation of the superior colliculus neurons improved perceptual task performance when visual stimuli appeared at locations encoded by the stimulated collicular neurons. Furthermore, Zénon and Krauzlis (2012) reported a perception deficit for stimuli presented at a location encoded by visual collicular neurons that were previously inactivated, but not for distracting stimuli presented outside the inactivated collicular receptive field. More recently, Bogadhi et al. (2020) have shown that superior colliculus inactivation modulates neural correlates of high-level visual functions (e.g. spatial and object-selective attention, stimulus detection) on the superior temporal sulcus in monkeys. Overall, these findings suggest that the superior colliculus can bias the cortical mechanisms of stimulus detection and selection. Further, Fecteau et al. (2004) showed an increase of target-related collicular response and a corresponding reduction of target-directed saccade onset time when the target was validly cued by another stimulus appearing at the same location ~50ms in advance. A 50ms CTOA is arguably sufficient time for bottom-up collicular modulation of target processing in primary visual cortex, but this mechanism seems less plausible for the ~24ms CTOA and low-contrast cues of our second experiment.

We propose that the cue-induced SLR modifications reported here reflect a spatiotemporal integration of the low-contrast and high-contrast stimuli accomplished subcortically through the tecto-reticolo-spinal circuits, rather than via cortical top-down feedback mechanisms. More specifically, we propose that the express visuomotor response in the valid cue conditions was faster than control because it was superimposed upon residual activity in the superior colliculus originating from the low-contrast cue. Functionally, this might aid the onset of rapid visuomotor responses to visual stimuli spatially congruent with weak sensory events that were recently experienced.

### Methodological considerations and future directions

Cueing the target location modified both volitional and express visuomotor responses, which may reflect priming mechanisms of top-down origin for the symbolic cues and bottom-up origin for low-contrast cues. However, it is unclear which cue type had the highest saliency to modulate the SLR expression, at least for the cue paradigms adopted here. Future studies should use different versions of our cueing paradigms to further delineate the neural mechanisms behind this express visuomotor behaviour in humans.

In this study, we reasoned that the effects of the symbolic cue reflected a cortical top-down priming of visuomotor networks, including the putative subcortical SLR-network. However, alternative interpretations might explain our observations. In the control conditions, the target appeared randomly to the left or right of participants’ dominant hand. Therefore, two distinct and competing motor programs could be prepared and coexist in the subcortical circuitry until that compatible with the actual target location was chosen and released. The integration between visual and motor-preparation signals could be facilitated if the competition between prepared motor programs is resolved, at least partially, before the stimulus presentation by cueing the target location. This would be expected to potentiate the SLR expression when the stimulus appears at a location congruent with the cue-related motor program and impair it when the prepared motor program mismatches the target location. For example, visual inputs to the superior colliculus might quickly trigger the nodes that are involved in the release of prepared responses (e.g. brainstem reticular formation nuclei; see for review Marinovic and Tresilian 2016; Carlsen and Maslovat 2019). Noteworthy, these hypotheses are consistent with the positive and negative cue-induced SLR gains observed in the first experiment. However, motor preparation mechanisms cannot underlie the effects of low-contrast cues because they were barely detectable, had weak predictive value (50%) and appeared too shortly (~24ms) before the high-contrast target to allow the pre-target preparation of a specific motor response. Nonetheless, we acknowledge that neural mechanisms consistent with motor preparation might contribute to SLR generation and, therefore, should receive attention for future investigations on this express visuomotor behaviour.

### Conclusions

This study has shown that cueing the location of a visual target modulates express visuomotor responses in humans. Symbolic cues appear able to modify express visuomotor behaviour via cortical top-down feedback signals to the putative subcortical SLR-network, including the superior colliculus and its downstream reticulo-spinal circuits. These phenomena illustrate a mechanism by which cognitive expectations can modulate the critical nodes for SLR generation to speed-up the visuomotor responses to expected visual events. By contrast, the effects of low-contrast cues appear to reflect exogenous priming mechanisms, potentially evolving subcortically via the superior colliculus. These mechanisms might aid the spatiotemporal integration of spatially congruent visual signals along the tecto-reticulo-spinal pathway and facilitate rapid response initiation when a salient stimulus follows a weak visual event. Overall, our findings help to constrain models of the neural mechanisms responsible for express visuomotor responses in humans.

## Acknowledgements

This work was supported by operating grants from the Australian Research Council (DP170101500) awarded to T.J. Carroll, B.D. Corneil, G.E. Loeb and G. Wallis.

## REFERENCES

Atsma J, Maij F, Gu C, Medendorp WP, Corneil BD. Active braking of whole-arm reaching movements provides single-trial neuromuscular measures of movement cancellation. J Neurosci. 38:4367–4382, 2018.

Baluch F and Itti L. Mechanisms of top-down attention. Trends Neurosci. 34:210–224, 2011.

Basseville M and Nikiforov IV. Detection of abrupt changes: theory and application (Information and System Sciences Series). Prentice Hall. 1993.

Basso MA, May PJ. Circuits for action and cognition: A view from the superior colliculus. Annu Rev Vis Sci. 3:197–226, 2017.

Boehnke SE and Munoz DP. On the importance of the transient visual response in the superior colliculus. Curr Opin Neurobiol. 18:544–551, 2008.

Bogadhi AR, Katz LN, Bollimunta A, Leopold DA, Krauzlis RJ. Midbrain activity shapes high-level visual properties in the primate temporal cortex. Neuron. 2020 Dec 8:S0896-6273(20)30928-4. doi: 10.1016/j.neuron.2020.11.023. Epub ahead of print. PMID: 33338395.

Brainard DH. The Psychophysics Toolbox. Spat Vis. 10:433–436, 1997.

Carlsen AN, Maslovat D. Startle and the StartReact effect: physiological mechanisms. J Clin Neurophysiol. 36:452–459, 2019.

Carroll TJ, McNamee D, Ingram JN, Wolpert DM. Rapid visuomotor responses reflect value-based decisions. J Neurosci. 39:3906–3920, 2019.

Contemori S, Loeb GE, Corneil BD, Wallis G, Carroll TJ. The influence of temporal predictability on express visuomotor responses. J Neurophysiol. 2020 Dec 23. 10.1152/jn.00521.2020. Epub ahead of print. PMID: 33357166.

Corneil BD and Munoz DP. Overt responses during covert orienting. Neuron. 82, 1230–1243, 2014.

Corneil BD, Munoz DP, Chapman BB, Admans T, Cushing SL. Neuromuscular consequences of reflexive covert orienting. Nat Neurosci. 11:13–15, 2008.

Dash S, Peel TR, Lomber SG, Corneil BD. Frontal eye field inactivation reduces saccade preparation in the superior colliculus but does not alter how preparatory activity relates to saccades of a given latency. eNeuro. 5: ENEURO.0024–18.2018, 2018.

Dorris MC, Klein RM, Everling S, Munoz DP. Contribution of the primate superior colliculus to inhibition of return. J Cogn Neurosci. 14:1256–1263, 2002.

Dorris MC and Munoz DP. Saccadic probability influences motor preparationsignals and time to saccadic initiation. J Neurosci. 18:7015–7026, 1998.

Dorris MC, Olivier E, Munoz DP. Competitive integration of visual and preparatory signals in the superior colliculus during saccadic programming. The Journal of Neuroscience. 27(19), 5053–5062, 2007.

Dorris MC, Pare M, Munoz DP. Neuronal activity in monkey superior colliculus related to the initiation of saccadic eye movements. J Neurosci. 17:8566–8579, 1997.

Fecteau JH, Bell AH, Munoz DP. Neural correlates of the automatic and goal-driven biases in orienting spatial attention. J Neurophysiol. 92(3):1728–37, 2004.

Fiehler K, Brenner E, Spering M. Prediction in goal-directed action. J Vis. 10:1–21, 2019.

Fischer B. and Boch R. Saccadic eye-movements after extremely short reaction-times in the monkey. Brain Res. 260:21–26, 1983.

Gandhi NJ and Katnani HA. Motor functions of the superior colliculus. Annu Rev Neurosci. 34:205–231, 2011.

Gu C, Pruszynski JA, Gribble PL, Corneil BD. A rapid visuomotor response on the human upper limb is selectively influenced by implicit motor learning. J Neurophysiol. 121:85–95, 2019.

Gu C, Pruszynski JA, Gribble PL, Corneil BD. Done in 100 ms: Path-dependent visuomotor transformation in the human upper limb. J Neurophysiol. 119:1319–1328, 2018.

Gu C, Wood DK, Gribble PL, Corneil BD. A trial-by-trial window into sensorimotor transformations in the human motor periphery. J Neurosci. 36:8273–82, 2016.

Haith AM, Pakpoor J, Krakauer JW. Independence of movement preparation and movement initiation. J Neurosci. 36:3007–3015, 2016.

Kingdom, F. A. A., and Prins, N. (2016). “Chapter 5: Adaptive methods,” in Psychophysics (San Diego, CA: Academic Press), 119–148.

Klein, R.M. Inhibition of return. Trends Cogn Sci. 4, 138–147, 2000.

Knudsen EI. Neural circuits that mediate selective attention: a comparative perspective. Trends Neurosci. 41:789–805, 2018.

Kozak AR, Cecala AL, Corneil BD. An emerging target paradigm evokes fast visuomotor responses on human upper limb muscles. J Vis Exp. e61428, doi:10.3791/61428, 2020.

Kozak RA, Kreyenmeier P, Gu C, Johnston K, Corneil BD. Stimulus-locked responses on human upper limb muscles and corrective reaches are preferentially evoked by low spatial frequencies. eNeuro. 6(5):ENEURO.0301–19. 2019.

Marino RA, Levy R, Boehnke S, White BJ, Itti L, Munoz DP. Linking visual response properties in the superior colliculus to saccade behavior. Eur J Neurosci. 35:1738–1752, 2012.

Marinovic W, Tresilian JR. Triggering prepared actions by sudden sounds: reassessing the evidence for a single mechanism. Acta Physiol. 217:13–32, 2016.

Moore T and Armstrong K. Selective gating of visual signals by microstimulation of frontal cortex. Nature. 421, 370–373, 2003.

Muller J et al. Microstimulation of the superior colliculus focuses attention without moving the eyes. Proc. Natl. Acad. Sci.U.S.A. 102, 524–529, 2005.

Pare M and Munoz DP. Saccadic reaction time in the monkey: advanced preparation of oculomotor programs is primarily responsible for express saccade occurrence. J Neurophysiol. 76:3666–3681, 1996.

Peel TR, Dash S, Lomber SG, and Corneil BD. Frontal eye field inactivation diminishes superior colliculus activity, but delayed saccadic accumulation governs reaction time increases. J Neurosci. 37: 728 11715–11730, 2017.

Pelli DG. The VideoToolbox software for visual psychophysics: Transforming numbers into movies. Spat Vis. 10:437–442, 1997.

Posner MI. Orienting of attention: Then and now. Q J Exp Psychol (Hove). 69(10):1864–1875, 2016.

Pruszynski AJ, King GL, Boisse L, Scott SH, Flanagan RJ, Munoz DP. Stimulus-locked responses on human arm muscles reveal a rapid neural pathway linking visual input to arm motor output. Eur J Neurosci. 32:1049–1057, 2010.

Pruszynski JA, Kurtzer I, Scott SH. Rapid motor responses are appropriately tuned to the metrics of a visuospatial task. J Neurophysiol. 100:224–238, 2008.

van Ede F, de Lange FP, Maris E. Attentional cues affect accuracy and reaction time via different cognitive and neural processes. J Neurosci. 2012; 32:10408–10412.

Wood DK, Gu C, Corneil BD, Gribble PL, Goodale MA. Transient visual responses reset the phase of low-frequency oscillations in the skeletomotor periphery. Eur J Neurosci. 42:1919–1932, 2015.

Zenon A and Krauzlis RJ. Attention deficits without cortical neuronal deficits. Nature. 489, 434–437, 2012.

